# Endogenous protection from ischemic brain injury by preconditioned monocytes

**DOI:** 10.1101/276923

**Authors:** Lidia Garcia-Bonilla, David Brea, Corinne Benakis, Diane Lane, Michelle Murphy, Jamie Moore, Gianfranco Racchumi, Xinran Jiang, Costantino Iadecola, Josef Anrather

## Abstract

Exposure to low dose lipopolysaccharide prior to cerebral ischemia is neuroprotective in stroke models, a phenomenon termed preconditioning. While it is well established that lipopolysaccharide-preconditioning induces central and peripheral immune responses, the cellular mechanisms modulating ischemic injury remain unclear. Here, we investigated the role of immune cells in the brain protection afforded by preconditioning and we tested whether monocytes may be reprogrammed by *ex vivo* lipopolysaccharide exposure thus modulating the inflammatory injury after cerebral ischemia in male mice. We found that systemic injection of low-dose lipopolysaccharide induces a distinct subclass of CD115^+^Ly6C^hi^ monocytes that protect the brain after transient middle cerebral artery occlusion in mice. Remarkably, adoptive transfer of monocytes isolated from preconditioned mice into naïve mice 7 hours after transient middle cerebral artery occlusion reduced brain injury. Gene expression and functional studies showed that IL-10, iNOS and CCR2 in monocytes are essential for the neuroprotection. This protective activity was elicited even if mouse or human monocytes were exposed *ex vivo* to lipopolysaccharide and then injected into male mice after stroke. Cell tracking studies showed that protective monocytes are mobilized from the spleen and reach brain and meninges, wherein they suppressed post-ischemic inflammation and neutrophils influx into the brain parenchyma. Our findings unveil a previously unrecognized subpopulation of splenic monocytes capable to protect the brain with an extended therapeutic window, and provide the rationale for cell therapies based on the delivery of autologous or allogeneic protective monocytes into patients with ischemic stroke.

**Significance Statement:** Inflammation is a key component of the pathophysiology of the brain in stroke, a leading cause of death and disability with limited therapeutic options. Here, we investigate endogenous mechanisms of protection against cerebral ischemia. Using LPS preconditioning as an approach to induce ischemic tolerance in mice, we found the generation of neuroprotective monocytes within the spleen from where they traffic to the brain and meninges suppressing post-ischemic inflammation. Importantly, systemic LPS preconditioning can be mimicked by adoptive transfer of *in vitro*-preconditioned mouse or human monocytes at translational relevant time points after stroke. This model of neuroprotection may facilitate clinical efforts to increase the efficacy of bone marrow mononuclear cell treatments in acute neurological diseases such as cerebral ischemia.

## Introduction

Induction of ischemic tolerance by the exposure to sub-injurious stressors, also known as preconditioning (PC), is a powerful mechanism to evoke endogenous neuroprotective programs (Iadecola and Anrather, 2011). In the brain, PC can be achieved by a variety of agents and stressors including ischemia, inflammatory mediators, metabolic blockers, anesthetics, cortical spreading depression, and seizures (Kirino, 2002). The immune system plays a key role in the establishment of ischemic tolerance. For example, toll like receptors (TLR), key innate immunity receptors, are potent mediators of cerebral PC (Pradillo et al., 2009; Marsh et al., 2009b; Wang et al., 2010; Garcia-Bonilla et al., 2014a), and activation of TLR4 by systemic administration of lipopolysaccharide (LPS) is a potent PC stimulus (Tasaki et al., 1997; Ahmed et al., 2000; Vartanian et al., 2011). However, circulating LPS is impermeable to the blood-brain barrier and does not gain access to the brain (Singh and Jiang, 2004; Banks and Robinson, 2010). Therefore, LPS is thought to induce ischemic tolerance by a systemic effect involving reprograming of the immune system (Smith et al., 2002), which, in turn, protects the brain by suppressing post-ischemic proinflammatory gene expression, endothelial and microglial activation, as well as leukocyte infiltration (Bauer et al., 2000; Rosenzweig et al., 2004; Lin et al., 2009; Marsh et al., 2009a; Vartanian et al., 2011). However, the cell type(s) on which systemic LPS acts to induce the widespread changes in the immune system underlying the development of cerebral PC remain to be defined.

Monocytes are bone marrow derived cells characterized by surface expression of CD11b and CD115, which are present in large quantities in the spleen (Swirski et al., 2009). In circulation they consist of functionally diverse subsets (Geissmann et al., 2003): Ly6C^+^CCR2^+^CX3CR1^low^ “inflammatory” monocytes, which contribute to the inflammatory process (Swirski et al., 2007; Tacke et al., 2007; Swirski et al., 2010), and Ly6C^−^CCR2^−^CX3CR1^high^ “resident” monocytes, which patrol the vasculature and participate in the resolution of inflammation (Auffray et al., 2007; Nahrendorf et al., 2007). A third subset, monocytic myeloid-derived suppressor cells (Serafini et al., 2004; Huang et al., 2006; Bronte, 2009), is present in the BM and share the same surface markers with inflammatory monocytes (Youn et al., 2008; Ostrand-Rosenberg, 2010; Forghani et al., 2012).

All monocytes subtypes have been implicated in ischemic injury in different organs, wherein they can play either protective or destructive roles (Arnold et al., 2007; Nahrendorf et al., 2007; 2010), but their involvement in ischemic PC remains unclear. Considering that monocytes express TLR4 and are a prime target of LPS (Muzio et al., 2000; Biswas and Lopez-Collazo, 2009) it is conceivable that they participate in LPS PC. Indeed, LPS administration induces BM egress of selected monocyte subtypes, such as Ly6C^+^ monocytes, which have the potential of modulating systemic immune responses (Serbina et al., 2008; Shi et al., 2011). Here, we have investigated the role of peripheral immune cells in the neuroprotection observed after LPS PC in mice. We found that LPS treatment induces a unique subpopulation of splenic monocytes endowed with a powerful neuroprotective capacity against experimental ischemic brain injury, mediated through suppression of meningeal inflammation. The findings highlight a previously unrecognized neuroprotective potential of a distinct monocyte subset and raise the prospect of novel cell-based stroke therapies based on monocyte-induced immunomodulation.

## Materials and Methods

### Mice

All procedures were approved by the institutional animal care and use committee of Weill Cornell Medicine and were performed in accordance with the ARRIVE guidelines (Kilkenny et al., 2011). Experiments were performed in 7-8 week-old wild type (JAX 000664), iNOS^−/−^ (MacMicking et al., 1995), IL10^−/−^ (JAX 002251, B6.129P2-Il10tm1Cgn/J), CCR2^RFP/RFP^ (JAX 017586, B6.129(Cg)-Ccr2tm2.1Ifc/J), Cox2^−/−^ (Morham et al., 1995), or GFP-transgenic (JAX 006567, C57BL/6-Tg(CAG-EGFP)131Osb/LeySopJ) male mice on a C57Bl/6J background obtained from Jackson Laboratory (JAX; Bar Harbor, ME) and from in-house colonies (iNOS^−/−^ and Cox2^−/−^ mice).

### Middle Cerebral Artery occlusion

Transient focal cerebral ischemia was induced using the intraluminal filament model of middle cerebral artery occlusion (MCAo), as described previously (Jackman et al., 2011). Occlusion and reperfusion was confirmed by measuring the cerebral blood flow (CBF) in the MCA territory by transcranial laser Doppler flowmetry. Only animals with CBF reduction of >85% during MCAo and CBF recovered by >80% after 10 minutes of reperfusion were included in the study (Jackman et al., 2011). Rectal temperature was monitored and kept constant (37.0 ± 0.5°C) during the surgical procedure and in the recovery period until the animals regained full consciousness. Topical lidocaine and bupivacaine were used for pre-operative analgesia and buprenorphine as post-operative analgesia.

### Measurement of infarct volume and neurological deficit

As described in detail elsewhere (Jackman et al., 2011), blinded assessment of infarct volume, corrected for swelling, was quantified 3 days after ischemia using Nissl stain on 30-µm-thick coronal brain sections and image analysis software (MCID, Imaging Research Inc., Ontario, Canada).

Sensorimotor deficit was assessed 1 day before and 3 days after MCAo by the tape test by an investigator blinded to the treatment groups (Benakis et al., 2016). Mice were tested at the start of the dark phase of the circadian day-night cycle. Briefly, an adhesive tape was placed on the dorsal aspect of both forepaws. Latency of tape contact with the mouth and tape removal was scored on videos recorded over 180 seconds. In the test performed 1 day before MCAo, mice received three trials with 10-minutes rest cycles in between, and on day 3 post-MCAo, mice received two trials. Results were expressed as time average of tape-contact or tape-removal latency in each test for the contralesional and lesional side.

### Intra-splenic injection of fluorescent latex beads

Intra-abdominal surgery was conducted to inject 50 µl (3.64x10^10^ particles/ml) of Fluoresbrite® Yellow Green Microspheres (0.50 µm; Polysciences Inc., Warrington, PA) into the mouse spleen capsule 6 hours after saline or LPS treatment. Meloxicam and buprenorphine were given for pre-emptive analgesia.

### LPS in vivo preconditioning

LPS was repurified from a commercial LPS preparation (*Salmonella enterica* serotype typhimurium; Sigma; L7261) according to a previously published protocol (Manthey and Vogel, 1994). Mice were injected with either purified LPS (0.5 mg/kg, i.p.) or saline (100 µl i.p.) 24 hours before MCAo.

### Cell isolation

Mice were anesthetized with pentobarbital and transcardially perfused with heparinized PBS. Meninges and spleen were dissected, placed on a 70-µm cell-strainer and gently homogenized in PBS. Brain cell isolation was carried out by mechanical method or enzymatic digestion with liberase when cell sorting was performed, as previously described (Garcia-Bonilla et al., 2014b; Benakis et al., 2016). Briefly, for enzymatic brain cell isolation, brain hemispheres were separated from the cerebellum and olfactory bulb and gently triturated in Hepes-HBSS buffer (138 mM NaCl, 5 mM KCl, 0.4 mM Na_2_HPO_4_, 0.4 mM KH_2_PO_4_, 5 mM D-Glucose, 10 mM Hepes) using a Gentle MACS dissociator (Miltenyi Biotec, Auburn, CA) following the manufacturer’s instructions. The suspension was digested with 62.5 µg/ml Liberase DH (Roche Diagnostics, Indianapolis, IN) and 50 U/ml DNAse I at 37°C for 45 minutes in an orbital shaker at 100 rpm. For mechanical cell isolation, brain hemispheres were homogenized in RPMI 1640 medium (Mediatech, Manassas, VA) using a dounce homogenizer. Brain cells isolated from enzymatic or mechanical procedures were washed and subjected to discontinuous 70/30% Percoll (GE Healthcare, Piscataway, NJ) density gradient centrifugation. Enriched-mononuclear cells were collected from the interphase. BM cells were flushed out from the femur and tibias and filtered through a 40-µm cell-trainer. Blood was drained from the submandibular vein into tubes containing sodium heparin and erythrocytes were lysed. Cell suspensions were subsequently stained for flow cytometric analysis.

### Flow cytometric analysis

Cells were stained using rat monoclonal antibodies (Table 1) and analyzed on a MACSQuant Analyzer 10 (Miltenyi Biotec). Brain infiltrating leukocytes were identified as CD45^hi^ and further gated by CD11b and CD11c markers. CD45^hi^CD11b^−^CD11c^+^ were identified as dentritic cells (DC). From the CD45^hi^CD11b^+^CD11c^−^ gate, both monocytes and neutrophils were identified as CD45^hi^CD11b^+^CD115^+^ and CD45^hi^CD11b^+^Ly6G^+^, respectively. Monocytes were further separated as Ly6C^hi^ or Ly6C^lo^. Infiltrating macrophages were classified as CD45^hi^CD11b^+^CD11c^−^CD115^−^Ly6G^−^. The same gating strategy was used to phenotype leukocyte in the meninges and peripheral organs. Appropriate isotype controls, ‘fluorescence minus one’ staining, and staining of negative populations were used to establish sorting parameters. Absolute cell numbers and frequencies were recorded. Samples were acquired and analyzed by an investigator blinded to the treatment groups.

**Table 1.** Antibodies used for cytometric analysis and sorting.

| **Antigen** | **Label** | **Clone** | **Isotype** | **Supplier** |
| --- | --- | --- | --- | --- |
| **CD16/CD32** | | 93 | rat IgG2b, $\lambda$ | Biolegend |
| **CD45** | BV 510 | 30F-11 | rat IgG2b | Biolegend |
| **CD11b** | APC/Cy7 | M1/70 | rat IgG2b, $\kappa$ | Biolegend |
| **CD11c** | PE/Cy7 | N418 | armenian<br>hamster IgG | Biolegend |
| **Ly6G** | PerCP/Cy5.5 | 1A8 | rat IgG2a, $\kappa$ | Biolegend |
| **CD115** | PE | AFS98 | rat IgG2a, $\kappa$ | Biolegend |
| **CD4** | F488 | RM4-5 | rat IgG2a, $\kappa$ | Biolegend |
| **TCR $\beta$** | PerCP/Cy5.5 | H57-597 | Armenian<br>hamster | Biolegend |
| **CD8a** | PE/Cy7 | 53-6.7 | rat IgG2a, $\kappa$ | Biolegend |
| **TCR $\gamma\delta$** | APC | GL3 | Armenian<br>hamster | Biolegend |
| **CD19** | APC/Cy7 | 6D5 | rat IgG2a, $\kappa$ | Biolegend |
| **CCR2** | APC | 475301 | rat IgG2b | R&D systems |

Antibody cocktail for monocyte purification by immunodepletion.

**Antibody cocktail for monocyte purification by immunodepletion.**
| **Antigen** | **Label** | **Clone** | **Isotype** | **Supplier** |
| --- | --- | --- | --- | --- |
| **CD117** | biotin | 2B8 | rat IgG2b, $\kappa$ | Biolegend |
| **CD45R**<br>**(B220)** | biotin | RA3-6B2 | rat IgG2a, $\kappa$ | Biolegend |
| **CD49** | biotin | HMa2 | armenian<br>hamster IgG | eBioscience |
| **TER 119** | biotin | TER-119 | rat IgG2b, $\kappa$ | Biolegend |
| **CD5** | biotin | 3 7.3 | rat IgG2a, κ | Biolegend |
| **Ly6G** | biotin | 1A8 | rat IgG2a | Biolegend |

Antibody cocktail for neutrophil purification by immunodepletion.

**Antibody cocktail for neutrophil purification by immunodepletion.**
| **Antigen** | **Label** | **Clone** | **Isotype** | **Supplier** |
| --- | --- | --- | --- | --- |
| **CD117** | biotin | 2B8 | rat IgG2b, κ | Biolegend |
| **CD45R**<br>**(B220)** | biotin | RA3-6B2 | rat IgG2a, κ | Biolegend |
| **CD49** | biotin | HMa2 | armenian<br>hamster IgG | eBioscience |
| **TER 119** | biotin | TER-119 | rat IgG2b, κ | Biolegend |
| **CD5** | biotin | 3 7.3 | rat IgG2a, κ | Biolegend |
| **CD115** | biotin | AFS98 | rat IgG2a | Biolegend |
| **F4/80** | biotin | BM8 | rat IgG2a, κ | Biolegend |

### Fluorescence Activated Cell Sorting

Isolated brain cells by the enzymatic procedure were stained for CD45, CD115, Ly6G and Ly6C as described above. Leukocytes isolated from brains of mice injected with LPS or saline that did not undergo MCAo, were identified as CD45^hi^ and further gated by CD115, and Ly6G and Ly6C markers. “Inflammatory” or “resident” monocytes, identified as CD45^hi^CD115^+^Ly6G^−^Ly6C^hi^ or Ly6C^lo^, respectively, were sorted on a FACSVantage cytometer (BD Bioscience) and collected in Trizol LS (Invitrogen, Carlsbad, CA) for transcriptomic profiling. Brains from three to four mice were pooled.

### Monocytes and neutrophil isolation by immunodepletion from BM

BM was flushed out from femurs and tibias and cells were dispersed by aspiration through a 20-gauge needle. After erythrolysis, BM cells were resuspended in MACS buffer (PBS supplemented with 2% FBS, 2 mM EDTA; 100µl/10^7^ cells). Cells were incubated with a biotinylated-antibody cocktail (see Table 1) and purified with anti-biotin-microbeads according to the manufacturers instructions (Miltenyi Biotech). Afterwards, cells were washed, resuspended in sterile physiological saline (5x10^6^ cells/ml) and administered by retroorbital injection (iv) in anesthetized mice. Cell preparations contained >90% CD115^+^Ly6C^hi^ monocytes as determined by flow cytometry analysis. Neutrophils were also purified from mouse BM of LPS-preconditioned animals by immunomagnetic negative enrichment essentially as described above with a different biotinylated-antibody cocktail (Table 1). Cell preparations contained >90% neutrophils as determined by flow cytometry (anti-Ly6G-PerCP-Cy5.5, clone 1A8, Biolegend).

### Generation of bone marrow-derived macrophages and myeloid-derived suppressor cells

Generation of bone marrow-derived macrophages (BMDM), classically activated macrophages (CAM) or alternatively activated macrophages (AAM) was conducted as described elsewhere with modifications (Weischenfeldt and Porse, 2008). After erythrocyte lysis isolated BM cells were resuspended (1x10^6^ cells/ml) in BM-medium (DMEM/F12, 10% FBS, 20% L929-conditioned medium, 100U/ml Penicillin and 100 µg/ml Streptomycin) and cultured in 5% CO_2_ at 37°C for 7 days. Fresh medium was added every 2-3 days. Polarization to CAM was induced by 100 U/ml IFNγ (Peprotech) for 48 hours and 10 ng/ml LPS for the last 24 hours. AAM were obtained by incubating BMM with 10 ng/ml recombinant murine IL-4 (R&D Systems) for 24 hours. Fully differentiated adherent macrophages were washed twice with warm DMEM/F12, and collected by detaching them with a stream of ice-cold DMEM/F12 containing 10% FBS. Cells were collected by centrifugation and resuspended in sterile saline (5x10^5^ cells/100 µl). Monocytic myeloid-derived suppressor cells (MDSC) were generated according to a previously published protocol (Highfill et al., 2010). Briefly, isolated BM cells were cultured in MDSC-medium (DMEM, 10% FBS, 10 mM Hepes, 1.5 mM L-arginine, 1.5 mM L-asparagine, 1.5 mM L-glutamine and 0.05 mM β-mercaptoethanol) in the presence of recombinant murine granulocyte macrophage colony-stimulating factor (250 U/ml; Biolegend) and granulocyte colony-stimulating factor (100 ng/ml; Peprotech) for 4 days. Interleukin-13 (Biolegend) was added (80 ng/ml) on day 3. Cells were harvested at day 4 and MDSC were isolated from each culture by immunomagnetic positive selection using anti-CD11b-PE antibody, anti-PE magnetic microbeads and LS column separation (Miltenyi Biotec). The purity of isolated cell populations was found to be greater than 90% by flow cytometry. Suppressor function was tested for inhibition of splenic CD4^+^ T cell proliferation in the presence of anti-CD3 and anti-CD28 antibodies (Figure 4B).

### Transcriptomic profiling

Total RNA was extracted from sorted Ly6C^hi^ brain monocytes for analysis of gene expression by Illumina Mouse WG-6 v2.0 Expression BeadChips. Total RNA was amplified with TargetAmp-Nano Labeling Kit for Illumina Expression BeadChip (Epicentre, Madison, WI). All procedures were conducted according to the manufacturer’s instructions by the Genomics Shared Resource supported by Roswell Park Cancer Institute and National Cancer Institute grant P30CA016056. Data were checked for quality with Illumina’s GenomeStudio gene expression module (v1.9.0). Summary data were background subtracted and normalized by variance stabilization using the beadarray (version 2.20.2) (Dunning et al., 2007) and limma (version 3.26.8) (Ritchie et al., 2015) packages within the R statistical environment (version 3.2.3). Analysis of differential expression was conducted in limma for probe sets filtered by spot detection (P < 0.05) and difference in mean signal intensity between treatment groups (dif > 4). Significantly changed genes were subjected to analysis of canonical pathways curated by DAVID Functional Annotation Bioinformatics Microarray Analysis (Sherman et al., 2007). Heat maps were generated in GenePattern (Reich et al., 2006) using genes selected by the differential gene expression analysis. Expression data have been deposited for public access in the NCBI Gene Expression Omnibus (GEO) under accession no. GSE84327.

### Quantitative Real-Time PCR

Quantitative determination of gene expression was examined as described before (Garcia-Bonilla et al., 2015). Total RNA was extracted from ischemic hemispheres and meninges using Trizol reagent. RNeasy Plus Mini Kit (Qiagen) was used to extract RNA from BM cells. After, cDNA was synthetized with iScript reverse transcription supermix for RT-qPCR (BioRad). qRT-PCR was conducted with cDNA in duplicate reactions using the Maxima SYBR Green/ROX qPCR Master Mix (2X) (Thermo Scientific). The reactions were incubated at 50°C for 2 minutes and then at 95°C for 10 minutes. A polymerase chain reaction cycling protocol consisting of 15 seconds at 95°C and 1 minute at 60°C for 45 cycles was used for quantification. Primer sequences (Invitrogen) are described in Table 2. Data were expressed as relative fold change over sham-operated control mice calculated by the 2^−ΔΔCt^ method (Livak and Schmittgen, 2001).

**Table 2.**
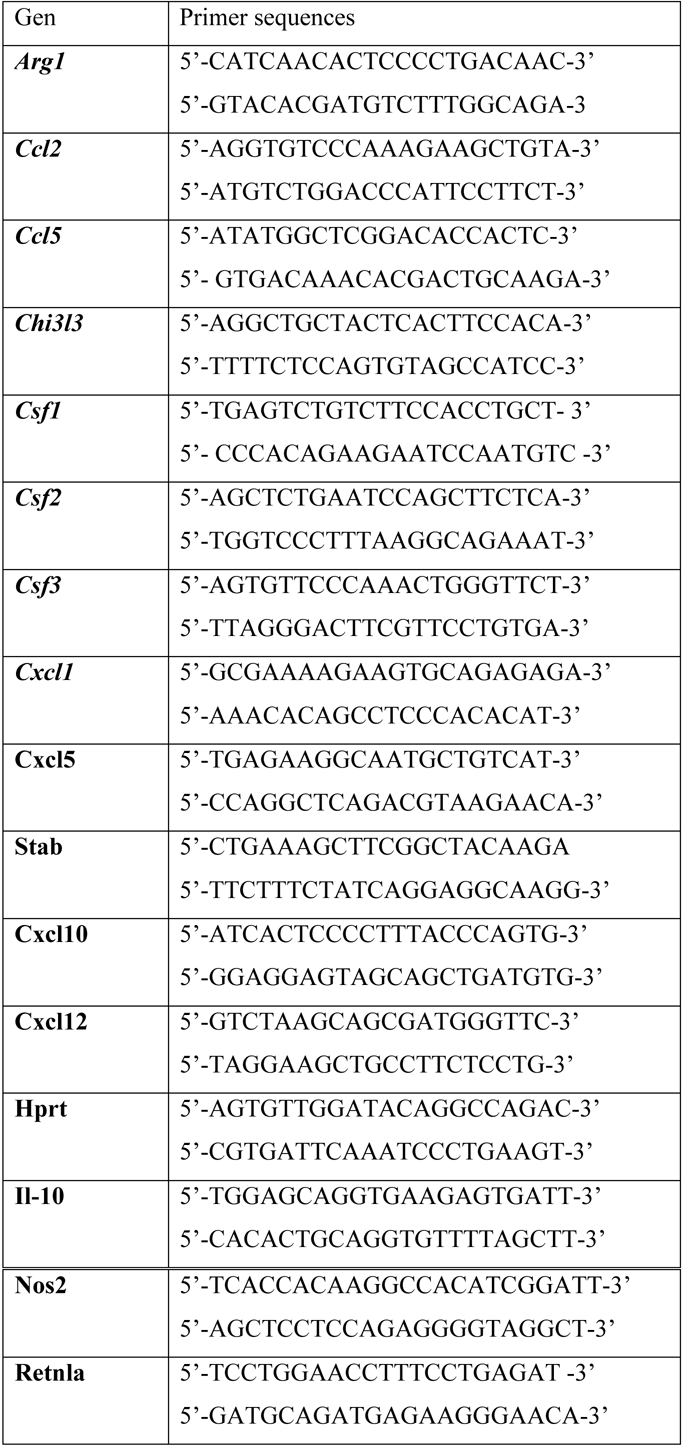
Primer sequences used for quantitative real-time PCR

### Histology

Histology of whole-mount meninges was performed as described (Louveau and Kipnis, 2015) with some modifications. Mice were anesthetized with sodium pentobarbital and perfused transcardially with ice-cold PBS followed by 4% paraformaldehyde (PFA) in PBS. Brains were extracted from the skull and the skullcap was post-fixed in 4% PFA-PBS overnight. After, the meninges (dura/arachnoid) were washed with PBS and dissected from the skull under a surgical microscope. Whole meninges were permeabilized with 0.5% Triton X-100 (Sigma) in PBS (PBST) for 30 minutes, blocked with 5% normal donkey serum (NDS) in 0.1% PBST for 1 hour and incubated with anti-Collagen IV (1:400, rabbit polyclonal; #ab6586, Abcam, RRID:AB_305584) antibody for 2 hours in 1%NDS-0.1% PBST. The meninges were washed with 0.1% PBST, incubated with Cy5-conjugated secondary antibody (1:200; Jackson ImmunoResearch Laboratories) and mounted on slices using ProLong® Gold Antifade mountant with DAPI (Thermo Fisher Scientific). All the staining procedure was performed at room temperature. Immunostaining was visualized by epi-fluorescent (Olympus IX83, Waltham, MA) or confocal microscopy (Leica TCS SP5, Mannheim, Germany).

Dissected brains and spleens were post-fixed in 4% PFA-PBS overnight at 4°C, immersed in 30% sucrose solution and frozen. Coronal brain sections and transverse sections of the spleen were cut (16 µm thickness) using a cryostat and mounted on slides for nuclear staining. Sections were permeabilized with 0.5%-PBST and then incubated with TO-PRO®-3 Iodide (Thermo Fisher Scientific) for 30 minutes at room temperature. Slices were washed with 0.1% PBST, mounted with FluorSave(tm) Reagent (Millipore) and visualized by confocal microscopy.

### Arginase and NOS activity

Arginase activity was assayed essentially as described previously (Corraliza *et al.*, 1994). Briefly, 1x10^5^ isolated BM monocytes from saline or LPS treated mice (24 hours) were lysed in 50 µl of 0.1% Triton X-100 containing pepstatin, aprotinin and antipain and incubated on a shaker for 30 minutes at room temperature. The enzyme was activated after adding 50 µl of 10 mM MnCl_2_, 50 mM Tris-HCl, pH 7.5 at 55°C for 10 minutes. After addition of equal volume of 0.5 M arginine, pH 9.7, arginine hydrolysis was performed at 37°C for 60 minutes. Reactions were stopped by acidification and the urea formed was colorimetrically quantified at 540 nm after addition of α-isonitrosopropiophenone and heating at 100°C for 45 minutes. NOS activity was assessed by measuring nitrite production. Briefly, isolated BM monocytes were resuspended in RPMI 1640 supplemented with 10% FBS and penicillin/streptomycin at a density of 5x10^5^ cells/ml and re-stimulated with 100 U/ml IFNγ and 100 ng/ml LPS for 24 hours at 37°C in a humidified atmosphere containing 5% CO_2_. Nitrite in supernatants was determined by the Griess reaction according to the manufacturers suggestions (Promega; #g2930).

### IL-10 protein level determination

IL-10 production in isolated BM monocytes was measured by ELISA (Thermo Fisher ScientificCat# BMS614/2, RRID:AB_2575685). Briefly, isolated BM monocytes were incubated in RPMI 1640 supplemented with 10% FBS and penicillin/streptomycin and re-stimulated with 100 ng/ml LPS for 16 hours at 37°C in a humidified atmosphere containing 5% CO_2_. Secreted IL-10 was assayed in cell culture supernatants.

### Experimental Design and Statistical Analysis Adoptive transfer of *in vivo*-preconditioned monocytes

A first study was performed to test whether the transfer of isolated monocytes from LPS-PC mice induces neuroprotection in mice that undergo MCAo. To this end, monocytes were purified by immunodepletion from BM cells of 7-8 week-old wild type, which were treated with saline or LPS (0.5 mg/kg in 100 µl saline i.p.) 24 hours before. A different researcher concealed to the surgical procedures, injected the isolated monocytes (5x10^5^ cells in 100 µl of sterile saline/mouse; i.v) into mice at different time points before or after MCAo. The mice were randomly allocated to receive either LPS- or Saline-monocytes. The infarct volume was analyzed 72 hours after MCAo to assess brain damage and compared to mice that received either saline or LPS (i.p) or that received adoptively transferred neutrophils (5x10^5^ cells in 100 µl of sterile saline/mouse; i.v).

A second study was conducted to test whether iNOS, IL-10, CCR2 or Cox2 molecules are important in conferring the protective effect of the monocytes. In these experiments, monocytes were purified by immunodepletion from BM cells of 7-8 week-old wild type, iNOS^−/−^, IL-10^−/−^, CCR2^RFP/RFP^ or Cox2^−/−^ mice, which were treated with saline or LPS (0.5 mg/kg in 100 µl saline i.p.) 24 hours before. The monocytes were injected (5x10^5^ cells in 100 µl of sterile saline/mouse; i.v) into mice 7 hours after MCAo. Infarct volumes were analyzed 72 hours after MCAo.

### Adoptive transfer of bone marrow-derived macrophages and myeloid-derived suppressor cells

A third study was conducted to assess whether bone marrow-derived macrophages that underwent classical (CAM) or alternative (AAM) activation, and myeloid-derived suppressor cells, have also the capacity of neuro-protect mice undergoing MCAo when adoptively transfer. Mice were randomly allocated to receive BMM, CAM, AAM or MDSC (5x10^5^ cells in 100 µl of sterile saline/mouse; i.v) 7 hours after MCAo. An investigator concealed to surgical procedures conducted macrophage transfers. Infarct volumes were analyzed 72 hours after MCAo.

### Adoptive transfer of *ex vivo*-preconditioned monocytes

In a forth study we tested the translational potential of transferring monocytes that are *ex vivo*-treated with LPS into mice undergoing MCAo. In these experiments, mouse monocytes isolated from BM of 7-

8 week-old naïve mice as described above, and human peripheral blood monocytes (79% CD14^+^CD16^low^ cells), purchased from Lifeline Cell Technology, were incubated with purified LPS (100 ng/ml) or vehicle (PBS) in RPMI with 10% FBS (2x10^6^ cells/ml) at 37°C and 5%CO_2_ for 2 hours. After, the monocytes were washed, resuspended in sterile saline and injected (5x10^5^ cells/100 µl saline; iv) into mice 7 hours after MCAo. Mice were randomly allocated to LPS- or PBS-*ex vivo* monocytes groups. An investigator blinded to surgical procedures and monocyte treatment performed monocyte transfers experiments. Infarct volumes were analyzed 72 hours after MCAo.

### Statistical Analysis

Data were analyzed using GraphPad Prism (Vers. 6) software. Intergroup differences were assessed for parametric data (Shapiro-Wilk normality test) by student’s t-test, one-way ANOVA or two-way ANOVA followed by post-hoc tests as appropriate. For non-parametric data, Mann-Whitney or Kruskal-Wallis test was used. Data are expressed as mean ± standard error of mean, and differences were considered statistically significant for a p value < 0.05 at a 95% confidence level.

## Results

### LPS preconditioning increases Ly6C^hi^ monocytes in brain and meninges

First, we sought to determine if administration of neuroprotective doses of LPS (Tasaki et al., 1997; Marsh and Stenzel-Poore, 2008) leads to an increase in immune cells in the CNS. LPS administration induced the recruitment of monocytes to the brain, identified by the surface markers CD115, when compared to mice receiving saline (Figure 1A, B). The increase was observed 24 hours after LPS and was sustained for at least 72 hours, a time window consistent with the protective effect afforded by LPS preconditioning (Rosenzweig et al., 2007). Although neutrophils increased 3.7 folds in blood 24 hours after LPS (Figure 1D), they were increased in brain only at 24 hours, an effect that did not reach statistical significance (Figure 1B). We then investigated whether the brain-associated monocytes were of the “inflammatory” (Ly6C^hi^) or “resident” (Ly6C^lo^) phenotype. CD115^+^Ly6C^hi^ monocytes were significantly higher in brains 24 hours after LPS compared to saline (Figure 1C). This was not correlated with an increase in the peripheral blood where CD115^+^Ly6C^hi^ cells after LPS was not different from saline controls (Figure 1C). In addition to the brain, CD115^+^ Ly6C^hi^ monocytes also increased in the meninges 24 hours after LPS treatment (Figure 1E), while no change in meningeal neutrophils was observed (Figure 1E). Non-myeloid immune cells in brain and meninges were not affected by LPS PC (data not shown). Therefore, LPS administration leads to accumulation of predominantly CD115^+^Ly6C^hi^ monocytes in brain and meninges.

**Figure 1.**
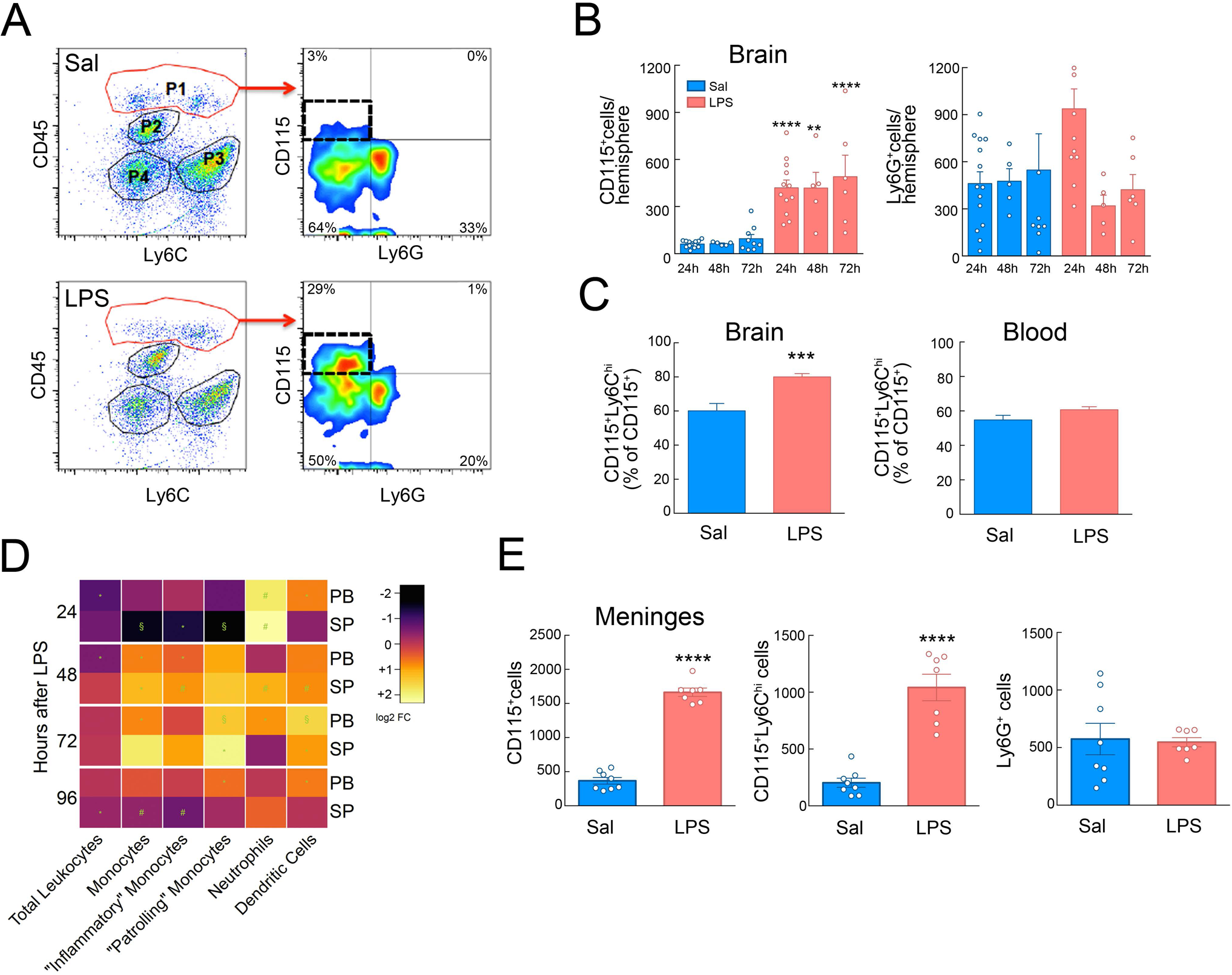
Systemic LPS preconditioning induces selective recruitment of inflammatory monocytes to brain and meninges. **A**. Flow cytometry analysis of isolated brain cells separated P1, P2, P3 and P4 populations based on CD45 and Ly6C expression. P1 identified infiltrating brain leukocytes (CD45^hi^) 24 hours after LPS injection (0.5 mg/kg; i.p.); P2 corresponded to microglia (CD45^int^Ly6C^−^); P3 identified endothelial cells (CD45^−^Ly6C^+^) and P4 corresponded to CD45^−^Ly6C^−^ cells. Further analysis of CD115 and Ly6G expression on infiltrating leukocytes (P1) identified recruited monocytes (CD45^hi^CD115^+^Ly6G^−^) and neutrophils (CD45^hi^CD115^−^Ly6G^+^). An increase of infiltrating monocytes was observed in LPS-treated mice as compared to saline-injected mice. **B**. CD115^+^ cells quantification in the brain indicated a sustained increase of monocytes in LPS-treated mice during the first 72 hours after LPS injection. A transient increase in the Ly6G^+^ cell number was observed at 24 hours in LPS injected mice, although statistical changes were not found in brain neutrophils between saline and LPS treatment over the first 72 hours (*n*=5-14 mice/group). Statistical analysis: two-way ANOVA (interaction, F (2, 44) = 0.07020, P = 0.9323; time, F (2, 44) = 0.4909, P = 0.6154; treatment, F (1, 44) = 55.76, P < 0.0001) followed by Sidak’s post-hoc test; **P < 0.05; ****P < 0.0001. **C.** Quantification of the percentage of brain and blood inflammatory monocytes (CD115^+^Ly6C^hi^) in saline (*n*=12) or LPS-treated (*n*=13) mice 24 hours after treatment. Increased Ly6C^hi^ percentage was observed in brain, but not in blood, of LPS-treated mice. Statistical analysis: Unpaired two-tailed Student’s t test (t=3.987, df=24, ***P=0.0005). **D**. Heatmap analysis of leukocyte numbers in peripheral blood (PB) or spleen (SP) from LPS-treated mice over saline-injected mice. Frequencies of leukocyte populations were calculated for each tissue at 24, 48, 72 or 96 hours after treatment. Log_2_ fold change was calculated for LPS over saline for each endpoint. Statistical analysis: Unpaired two-tailed Student’s t test; *P < 0.05; #P < 0.01; §P < 0.001. **E.** The total monocyte (CD115^+^) and inflammatory monocyte (CD115^+^Ly6C^hi^) number was increased in meninges of mice 24 hours after either saline (*n*= 8) or LPS (*n*= 7) injection. Neutrophils (Ly6G^+^) did not change. Statistical analysis: Unpaired two-tailed Student’s t test (t=16.93, df=13, ****P < 0.0001) for total CD115^+^ cells; Unpaired two-tailed Student’s t test (“t=7.207 df=13”, ****P < 0.0001) for CD115^+^Ly6C^hi^ cells.

### Adoptive transfer of monocytes from preconditioned donors protects naïve mice from ischemic brain injury

Having established that LPS preconditioning leads to recruitment of CD115^+^Ly6C^hi^ monocytes to brain and meninges, we investigated whether these cells contribute to the neuroprotection observed after LPS preconditioning. To this end, we isolated bone marrow monocytes 24 hours after LPS or saline administration and transferred them into naïve mice undergoing MCAo (Figure 2A). More than 90% of immunopurified BM monocytes were Ly6C^hi^ (Figure 2B). Adoptive transfer of monocytes (5x10^5^ cells iv) from LPS-trated mice 7 hours, but not 24 hours after MCAo resulted in a reduction in infarct volume similar to that of mice receiving LPS 24 hours before MCAo (Figure 2C, D). Adoptive transfer of monocytes 24 hours before MCAo resulted in partial protection that was not statistically significant (Figure 2D). Monocytes isolated from mice 24 hours after saline administration did not elicit neuroprotection when adoptively transferred to naïve mice 7 hours post-MCAo (Figure 2E). Similarly, adoptive transfer of neutrophils isolated from the BM of LPS-treated mice did not confer neuroprotection after adoptive transfer into naïve hosts 7 hours after MCAo (Figure 2E). These findings indicate that CD115^+^Ly6C^hi^ monocytes isolated from LPS-treated mice and transferred into naïve mice reproduce in full the protective effect of LPS PC.

**Figure 2.**
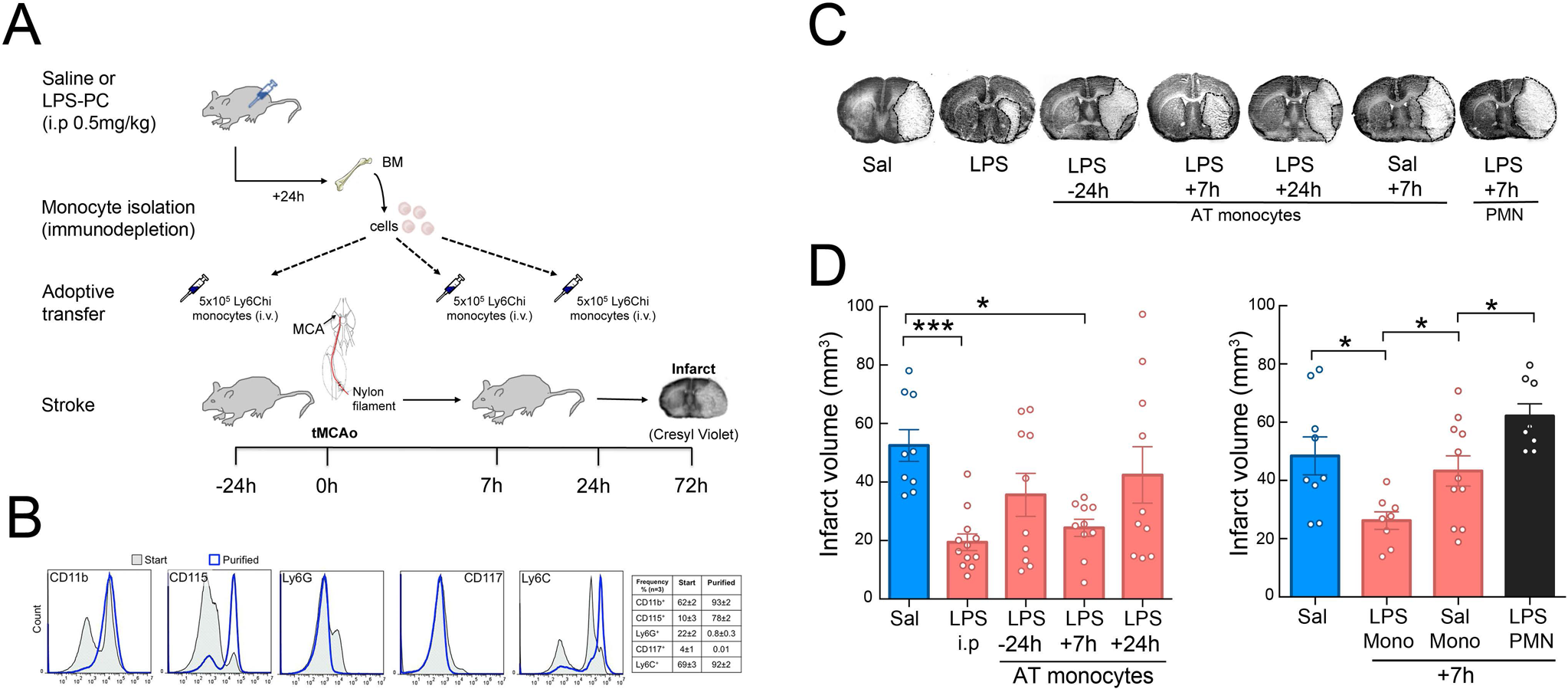
Adoptive transfer of LPS-primed monocytes isolated from preconditioned donors decreases infarct volume in stroke recipient mice. **A.** Experimental design for adoptive transfer of monocytes in stroke mice. Mice were injected with either LPS or saline and monocytes were isolated from bone marrows (BM) by immunodepletion 24 hours later. Saline or LPS-monocytes (5x10^5^ monocytes/100 µl) were injected intravenously into stroke mice 24 hours before transient MCAo or 7 or 24 hours post-MCAo. Infarct volume was determined by cresyl violet staining 72 hours after MCAo. **B**. Isolated bone marrow cells were stained for CD11b, CD115, Ly6G, CD117 and Ly6C markers before (start) and after (purified) immunodepletion. Stained cells were assayed by flow cytometry. Monocytes were isolated with high purity as evidenced by the expression of CD11b, CD115 and Ly6C markers and lack of expression of the neutrophil marker Ly6G and the hematopoietic stem cell marker CD117. **C**. Representative images of brain infarcts in saline-injected mice (Sal), LPS-preconditioned mice (LPS) or mice that received monocytes isolated from LPS-preconditioned mice 24 hours before MCAo (-24h) or 7 or 24 hours after MCAo (+7h and +24h, respectively). Mice that received monocytes from saline-injected mice are indicated as Sal +7h, and mice that received neutrophils isolated from LPS-injected mice are indicated as PMN +7h. The infarct (pale areas) is outlined in brain coronal sections stained with cresyl violet 72 hours after MCAo. AT stands for adoptive transfer. **D**. Infarct volume was significant smaller in either LPS preconditioned mice (LPS i.p.) or in mice receiving adoptively transferred LPS-monocytes at +7 hours (*n*= 9-12). Statistical analysis: Kruskal-Wallis (H = 15.65, P = 0.0035) followed by Dunn’s post-hoc test. *P < 0.05; ***P < 0.001. **E**. Adoptive transfer of monocytes isolated from saline-injected mice or neutrophils (PMN) from LPS-preconditioned mice at +7 hours post-MCAo did not reduce infarct volumes. (*n*= 8-11). Statistical analysis: one-way ANOVA F (3, 32) = 7.747, P=0.0006) followed by Holm-Sidak’s post-hoc test; *P < 0.05.

### Transcriptomic profiling of protective monocytes

Next, we examined if LPS induced brain-associated monocytes exhibit a gene expression profile different from those of saline-treated animals. Transcriptomic profiling of brain monocytes identified 270 genes that were up-regulated (> 4 folds) in monocytes isolated from brains of LPS-trated animals compared to monocytes of saline controls, while 414 were down-regulated (Figure 3A). Up-regulated genes were associated with immune and inflammatory responses, regulation of GTPase activity and cholesterol homeostasis, while down-regulated genes included members of cell adhesion, cell proliferation, positive transcriptional regulation, and angiogenesis pathways (Figure 3B). Genes associated with alternative macrophage polarization were highly upregulated, including *IL10* and *arginase 1* (Figure 3C). *IL10* and *Arg1* mRNA, assessed by RT-PCR, were also increased in monocytes isolated from the BM of LPS-treated mice (Figure 3D), in which a corresponding increase in IL-10 protein expression and arginase enzymatic activity was also observed (Figure 3E). Moreover, inducible nitric oxide synthase (*Nos2*) mRNA, a gene involved in LPS preconditioning (Kawano et al., 2007; Kunz et al., 2007), was also upregulated in BM monocytes isolated from LPS-treated mice (Figure 3D) and associated with increased NO production (Figure 3E). Other genes linked to alternatively activated macrophages like YM1 (*Chi3l1*), Fizz1/RELMα (*Retnla*), and stabilin 1 (*Stab1*) were also upregulated (Figure 3F). Therefore, the “protective” monocytes induced by LPS administration express selected genes typical of alternatively activated macrophages.

**Figure 3.**
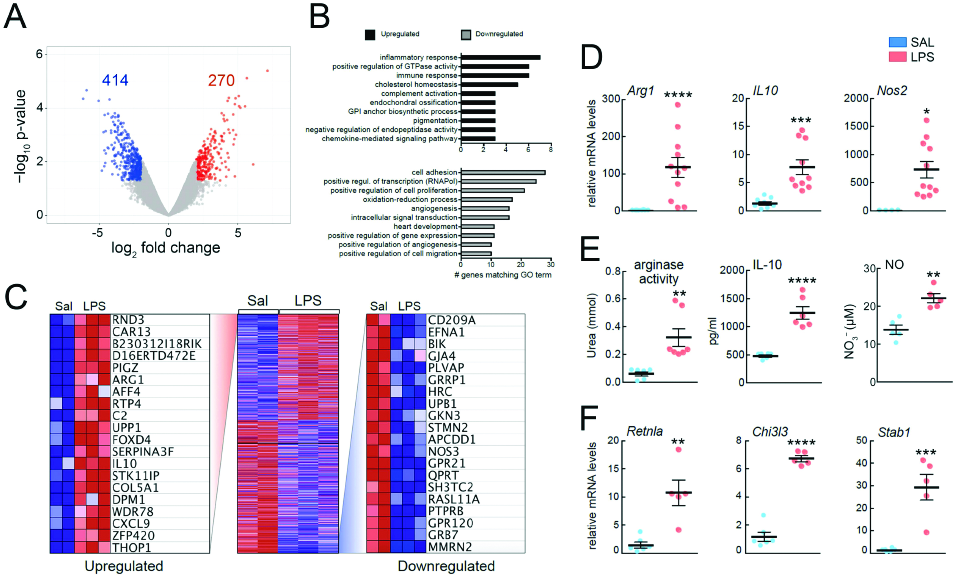
Transcriptomic analysis of brain monocytes from LPS preconditioned mice. **A**. Log_2_ fold-change vs. p-value plot of microarray data analysis of gene expression in isolated brain monocytes from LPS-preconditioned mice over saline-treated mice (N=2-3, *n*=8-12), highlighting down-regulated genes (blue, < 4 fold) and up-regulated genes (red, > 4 fold). **B**. Gene Ontology (GO) analysis of gene expression revealed up-regulation of genes associated with immune and inflammatory response, regulation of GTPase activity and cholesterol homeostasis and down-regulated genes members of cell adhesion, cell proliferation, positive transcriptional regulation, and angiogenesis pathways. **C**. Heatmap showing the top 20 genes upregulated and the top 20 genes downregulated in LPS vs. Sal-treated mice. **D.** Quantification of Arginase 1 (*Arg1*), IL-10 (*IL10*) and inducible nitric oxide gene (*Nos2)* expression by qRT-PCR in mouse bone marrow monocytes. Monocytes were isolated 24 hours after LPS or saline (SAL) injection. Data are expressed as relative n-fold changes over SAL group. *Arg1, IL10 and Nos2* mRNA were significantly upregulated in LPS-primed monocytes. Statistical analysis: *Arg1*, Mann Whitney test (U= 0, ****P< 0.0001); *IL-10*, Mann Whitney test (U= 0, ****P< 0.0001); *Nos2*, unpaired two-tailed Student’s t test (t=2.895, df=13,*P =0.0125). **E**. Quantification of arginase activity by determination of urea, IL-10 protein level by ELISA assay and iNOS activity by Griess reaction showed increased activities or protein level in BM monocytes of LPS-preconditioned mice compared to BM monocytes of saline-treated mice. Statistical analysis: urea, Mann Whitney test (U= 0, **P< 0.0012); IL-10, unpaired two-tailed Student’s t test (t=6.749, df=10, ****P < 0.0001); NO, unpaired two-tailed Student’s t test (t=4.83, df=8, **P=0.0013), respectively. **F**. Quantification of genes associated with alternatively activated macrophage polarization showed upregulated mRNA levels in BM monocytes isolated 24 hours after LPS-treatment over saline treatment. Statistical analysis: *Retnla*, unpaired two-tailed Student’s t test (t=4.371, df=9, **P=0.0018); *Chi3l3*, unpaired two-tailed Student’s t test (t=13.71, df=9, ****P < 0.0001), *Stab1*, unpaired two-tailed Student’s t test (t=5.426, df=9, ***P= 0.0004).

### Unlike monocytes, macrophages and myeloid-derived suppressor cells are not neuroprotective

Given that alternatively-activated macrophages have been observed in the brain after ischemia and could be beneficial due to their anti-inflammatory activity (Miró-Mur et al., 2015), we compared the neuroprotective ability of LPS-induced monocytes with that of bone marrow-derived macrophages that underwent classical (CAM) or alternative (AAM) activation. In addition, we assessed the protective capacity of myeloid-derived suppressor cells, a myeloid cell population with strong anti-inflammatory activity upregulated in the periphery after stroke (Serafini et al., 2004; Huang et al., 2006; Bronte, 2009; Liesz et al., 2015). To this end we generated CAM, AAM and MDSC from BMM *in vitro*, by exposing BMM to IFNγ/LPS, IL-4, or IL-4/IL-13 respectively (Weischenfeldt and Porse, 2008; Highfill et al., 2010). Successful polarization was confirmed by monitoring mRNA expression of genes associated with the respective subpopulations including *Nos2, Arg1, IL10, Nos2, Retnla, Chi3l3, Retnla*, and *Stab1* (Figure 4A). MDSC were tested for their ability to suppress CD4^+^ T cell proliferation (Figure 4B). All *in vitro* generated macrophage populations expressed CCR2, a chemokine receptor needed for macrophage entry into the brain and association with meninges (Figure 4C). Cells (5x10^5^) were then adoptively transferred to mice 7 hours after MCAo and infarct volumes determined 72 hours later. We found that none of the *in vitro* differentiated BMM populations reduced infarct volume after adoptive transfer (Figure 4D). Therefore, unlike LPS-induced protective monocytes, CAM, AAM and MDSC are unable to ameliorate ischemic brain injury.

**Figure 4.**
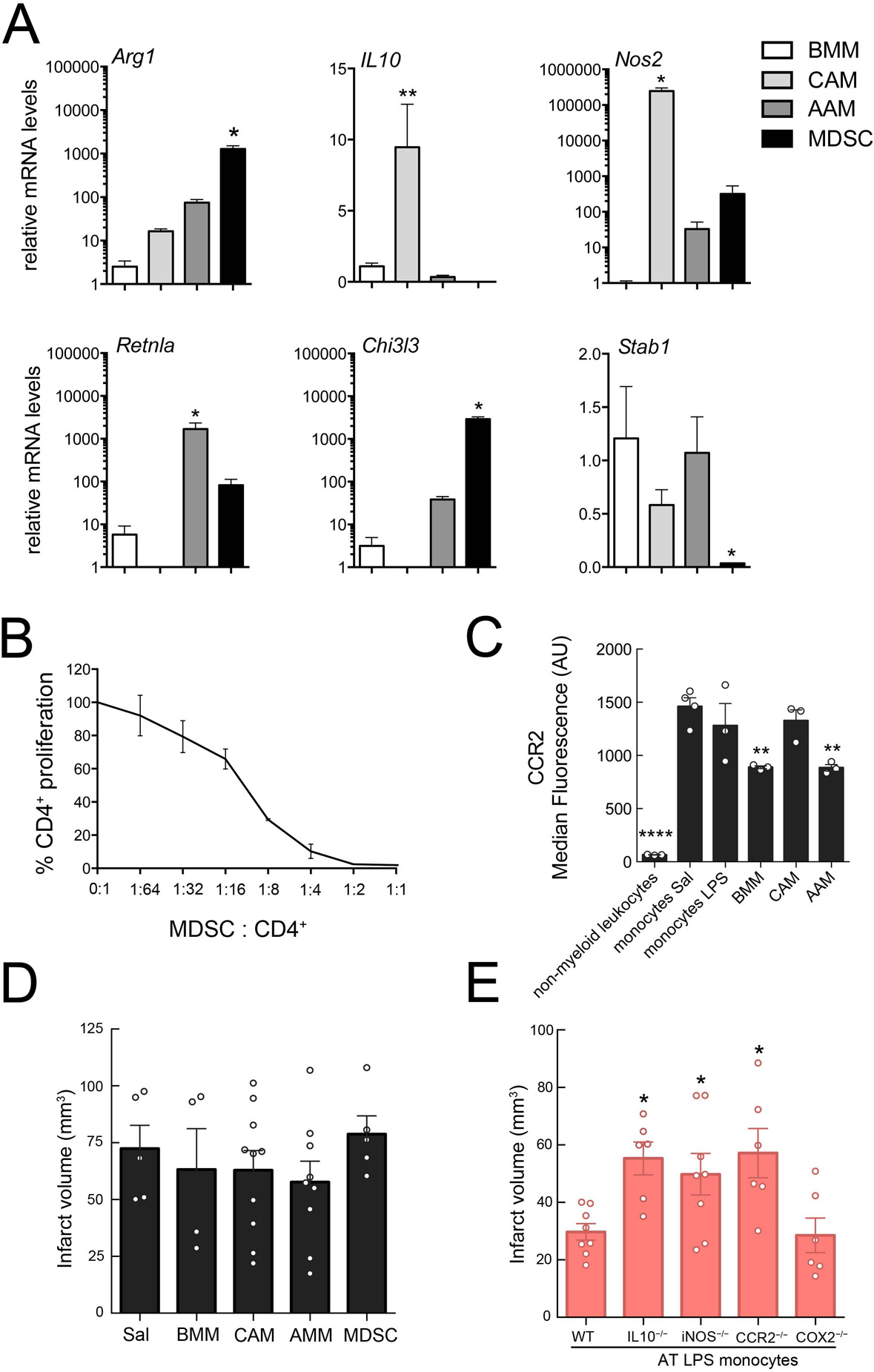
IL-10, iNOS and CCR2 are required for the neuroprotection induced by LPS-primed monocytes. **A**. Quantification of Arginase 1 (*Arg1*), IL-10 (*IL10*), iNOS (*Nos2*), *Retnla, Chi3l3* and stabilin 1 (*Stab1*) gene expression in cultured bone marrow-derived macrophages (BMM), classical or alternative activated bone marrow-derived macrophages (CAM and AAM, respectively), and myeloid-derived suppressor cells (MDSC) by qRT-PCR. Statistical analysis: Kruskal-Wallis followed by Dunn’s test vs. BMM; *Arg 1*, (H = 8.291, P = 0.0013), *P = 0.05; *IL10*, (H = 10.97, P < 0.0001), **P = 0.01; *Nos2*, (H = 9.859, P= 0.003), *P = 0.05; *Retnla*, (H = 9.929, P= 0.0012), *P = 0.05; *Chi3l3*, (H = 10.24, P= 0.006), *P = 0.05; *Stab1*, (H = 7.750, *P = 0.0286), *P = 0.05; n=3-4/group. **B**. Increment cells numbers of splenic CD4+ cells were stimulated with anti-CD3 and anti-CD28 antibodies in the presence of MDSC. Percentage of CD4+ T cell proliferation was calculated to assess suppressor function (n=2). X-axis values represent MDSC/splenic CD4+cell ratio. **C**. Quantification of the median fluorescence of CCR2 signal analyzed by flow cytometry of in non-myeloid leukocytes, bone marrow monocytes from saline (SAL) or LPS-treated mice, cultured bone marrow-derived macrophages (BMM) and classical or alternative activated bone marrow-derived macrophages (CAM and AAM, respectively). Statistical analysis: one-way ANOVA F (5, 13) = 26.13, P < 0.0001) followed by Holm-Sidak’s post-hoc test; **P < 0.01; ****P < 0.0001; n=3-4/group. **D.** The infarct volumes of mice receiving bone marrow-derived macrophages (BMM; +7 hours post-MCAo), BMM that underwent either classical (CAM) or alternative (AAM) activation, or BMM that were polarized to myeloid-derived suppressor cells (MDSC) were not different from those of mice treated with saline (Sal), (*n*=4-10 mice/group). Statistical analysis: Kruskal-Wallis (H = 2.833, P=0.7257). **E**. Mice receiving monocytes (7 hours after MCAo) isolated from IL-10^−^ /–, iNOS^−^/– or CCR2^−^/– LPS-preconditioned mice, but not from COX2^−^/– LPS-preconditioned mice, had larger infarcts as compared to the mice receiving monocytes isolated from wild type (WT) LPS-preconditioned mice (*n*=6-8 mice/group). AT stands for adoptive transfer. Statistical analysis: one-way ANOVA F (3, 32) = 4.175, P=0.0040) followed by Holm-Sidak’s post-hoc test; *P < 0.05.

### iNOS, IL-10 and CCR2 are required for the beneficial effect of monocytes

Because *Nos2* and *IL10* gene expression was upregulated in LPS-induced monocytes, we examined whether iNOS or IL-10 were required for their neuroprotective effects. To this end, BM monocytes isolated from iNOS^−/−^ or IL10^−/−^ LPS-treated mice were adoptively transferred (5x10^5^ cells) to wild type mice 7 hours after MCAo and infarct volume was determined 72 hours later. Neither iNOS^−/−^ nor IL10^−/−^ monocytes induced neuroprotection as compared to LPS-induced wild type monocytes (Figure 4E). Because CCR2 is required for the recruitment of “inflammatory” monocytes to sites of ischemic inflammation (Swirski et al., 2009; Garcia-Bonilla et al., 2016), we adoptively transferred monocytes from CCR2^−/−^ mice treated with LPS (Figure 4E). We found that CCR2^−/−^ monocytes failed to induce neuroprotection suggesting that monocyte migration to the brain and/or meninges is required for the protection. In contrast, LPS-induced monocytes isolated from mice deficient for the inflammatory enzyme cyclooxygenase-2 (COX2) were able to protect from ischemic brain injury, attesting to the specificity of iNOS, IL10, and CCR2 in the protective effect.

### Trafficking of protective monocytes to spleen, meninges and brain

Next, we investigated the potential source of the protective monocytes in mice undergoing LPS PC. Since we observed a significant depletion of splenic monocytes 24 hours after LPS PC while blood monocytes were not changed (Figure 1D), we tested whether monocytes are mobilized from the spleen and recruited to the brain and meninges. The spleen is a monocyte reservoir that can be mobilized after ischemia in other organs (Swirski et al., 2009), and splenic monocytes have been implicated in stroke pathology (Seifert et al., 2012; Dotson et al., 2014; Kim et al., 2014). To specifically label phagocytes in the spleen, fluorescent latex (Lx) beads (0.5 µm diameter), were injected into the spleen of mice 6 hours after they were treated with either saline or LPS. Eighteen hours later, histological examination of the spleen revealed that Lx beads distributed throughout the subcapsular sinus and the red pulp (Figure 5A). Flow cytometric analysis showed efficient uptake of Lx-beads by splenic CD115^+^Ly6G^−^ monocytes in LPS and saline treated animals (Figure 5B, C). However, Lx^+^ monocytes were found in blood, brain and meninges in LPS-treated animals, but not saline controls, indicating that LPS PC mobilizes splenic monocytes that are then recruited to brain and meninges (Figure 5B, C).

**Figure 5.**
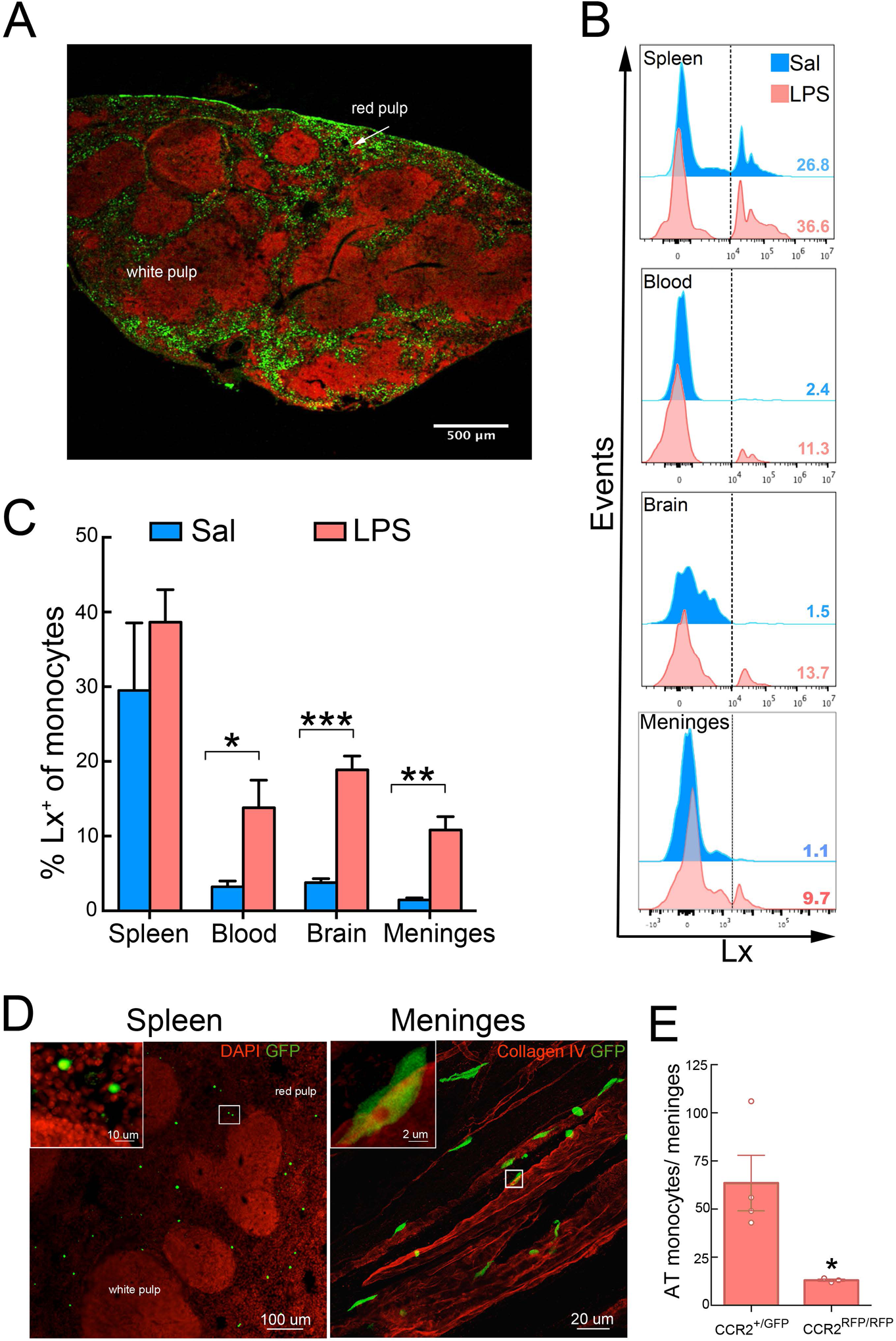
LPS induces trafficking of splenic monocytes to the brain and meninges. **A**. Representative image of the localization of fluorescent latex beads in the spleen 24 hours after intrasplenic injection (1.8x10^9^ particles). The beads (green) accumulated in the marginal zone and subcapsular red pulp. Fluorescent nuclear staining with the TO-PRO-3 (red) was used to reveal the structural anatomy of the spleen (transverse section). **B**. Representative flow cytometry histograms and quantification (**C**) of monocytes (CD115^+^ cells) showing the percentage of monocytes containing fluorescent latex beads (Lx) in the spleen, blood, brain and meninges 24 hours after either LPS-preconditioning or saline injection in mice (*n*=3-4/group). Lx were injected into the spleen 7 hours after saline or LPS treatment. Statistical analysis: unpaired two-tailed Student’s t test, blood (t=2.77, df=6, *P=0.0321), brain (t=7.81, df=6, ***P=0.0002) and meninges (t=5.22, df=6, **P=0.0019). **D**. Representative image of the spleen (left) and meninges (right) from mice that underwent 24 hours of MCAo and received GFP^+^-monocytes (5x10^5^ cells, iv) isolated from LPS-preconditioned mice at 7 hours post-MCAo. Transverse section of the spleen was stained with DAPI (red) for nuclear staining. Adoptively transferred monocytes accumulated in the red pulp of the spleen. In the meninges, monocytes were associated with meningeal blood vessel revealed by collagen IV (red) immunohistochemistry. **E**. Quantification of adoptively transferred monocytes (AT) in the meninges of wild type mice 24 hours after MCAo. Mice received either GFP^+^CCR2^+/+^ monocytes (CCR2^+/GFP^) or RFP^+^CCR2^−^/–-monocytes (CCR2^RFP/RFP^; 5x10^5^ cells, iv) isolated from GFP-wild type or CCR2^RFP/RFP^ LPS-preconditioned mice, respectively. GFP or RFP fluorescence was used to track transferred BM monocytes (*n*=4 mice/group). The genetic deletion of CCR2 resulted in a drop of meninges-associated monocytes. Statistical analysis: unpaired two-tailed Student’s t test, blood (t=2.96, df=5, *P=0.0315).

To gain an insight into the trafficking of adoptively transferred protective monocytes, we isolated BM monocytes from mice ubiquitously expressing green fluorescent protein (GFP) treated with LPS 24 hours earlier, and transferred them to wild type mice 7 hours after MCAo. Histological analysis of spleen, brain and meninges was performed 24 hours after MCAo. In contrast to monocytes in mice treated with LPS (Figure 5B, C), adoptively transferred GFP^+^ monocytes were found in the spleen and meninges but not the brain parenchyma (Figure 5D). The transfer to the meninges required CCR2 because CCR2^−^/– monocytes did not efficiently migrate to the meninges (Figure 5E). Therefore, spleen and meninges are major sites of monocytes localization both in LPS preconditioned mice and in mice adoptively transferred with monocytes from LPS-treated mice undergoing MCAo.

### LPS PC results in increased meningeal and decreased brain immune cell accumulation after stroke

We then examined whether the presence of PC-induced monocytes in brain and meninges has an impact on the post-ischemic infiltration of inflammatory cells in brain. Mice underwent LPS or saline treatment 24 hours before stroke and the cell populations in brain and meninges were assessed 48 hours after MCAo, a time point when peripheral immune cell infiltration is maximal (Garcia-Bonilla et al., 2014b; Benakis et al., 2016). While monocytes were increased in the meninges of LPS-PC mice compared to saline treated controls (Figure 6A), brain monocytes were unchanged (Figure 6B). In contrast, the number of neutrophils was not different in the meninges between groups, but brain-infiltrating neutrophils were reduced in LPS-PC mice (Figure 6A, B). CD4^+^ T cells, CD8^+^ T cells, γδT cells, B cells and NK cells trended lower without reaching statistical significance (data not shown). Together, these data indicate that PC-induced monocytes in the meninges are associated with reduced neutrophil trafficking into the brain after a stroke.

**Figure 6.**
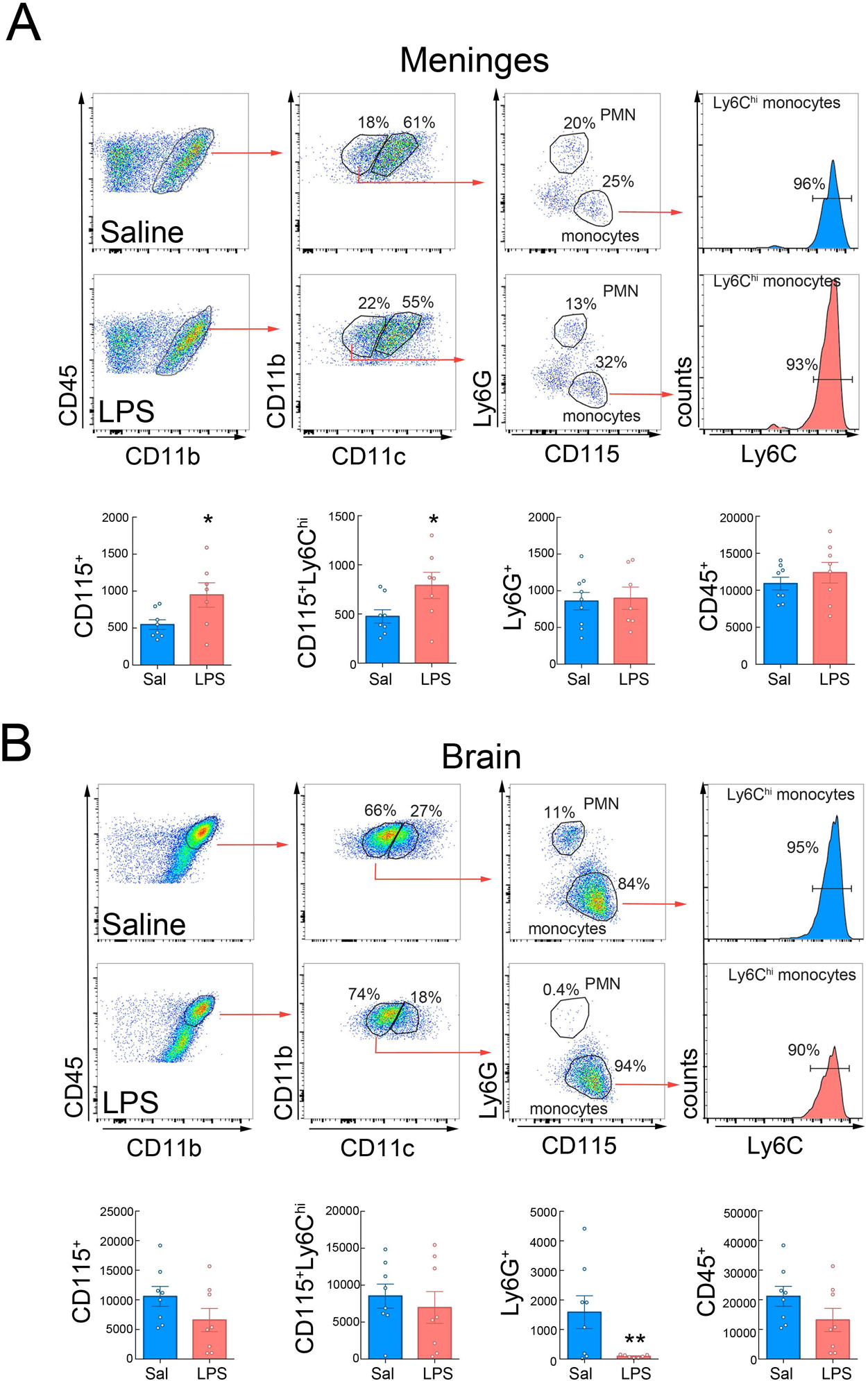
LPS preconditioning increases meningeal monocyte and decreases brain neutrophil recruitment after MCAo. Flow cytometry analysis of immune cell infiltration in the meninges (**A**) and the brain (**B**) of saline or LPS-preconditioned mice 48 hours after MCAo (*n*=8 mice/group). Flow cytometry plots depict gating strategy. Infiltrating leukocytes (CD45^hi^) were phenotyped as monocytes by high expression of CD11b and CD115 and low expression of CD11c and Ly6G (CD115^+^ cells). Inflammatory monocytes were subclassified by high Ly6C expression (CD115^+^Ly6C^hi^). Neutrophils (PMN) were identified by high expression of CD11b and Ly6G and low expression of CD11c and CD115 (Ly6G^+^ cells). **A**. Graph plots show increased inflammatory monocytes (CD115^+^ and CD115^+^Ly6C^hi^ cells) in the meninges of LPS-preconditioned mice following MCAo. Statistical analysis: CD115^+^, unpaired two-tailed Student’s t test, blood (t=2.381, df=13, *P=0.0332); CD115^+^Ly6C^hi^, unpaired two-tailed Student’s t test, blood (t=2.186, df=13, *P=0.0477); **B**. Graph plots show decreased brain neutrophils (PMN) in LPS-preconditioned mice following MCAo. Statistical analysis: Ly6G^+^, unpaired two-tailed Student’s t test, blood (t=2.502, df=13, *P=0.0265).

### *Ex vivo* treatment with LPS induces a neuroprotective phenotype in mouse and human monocytes

While our results are consistent with a direct effect of LPS on monocytes leading to a protective phenotype, indirect activation by down-stream effectors induced by LPS, such as pro-inflammatory cytokines, lipid mediators or DAMP, cannot be excluded. To address whether LPS acts directly on monocytes, we exposed monocytes isolated from naïve animals to LPS (100 ng/ml, 2 hours) or vehicle (PBS) *ex vivo*, before injecting them (5x10^5^ cell, iv) into mice 7 hours after MCAo. Mice receiving LPS-treated monocytes showed smaller infarcts (-55±10%; P<0.05) than mice receiving PBS treated monocytes (Figure 7A), a protective effect comparable to that observed in LPS-PC mice (Figure 2A and 7A). Importantly, the protection was lost when LPS-stimulated monocytes were lysed (Figure 7A). The reduction in brain injury was accompanied by a better neurological outcome, assessed by the tape test 72 hours post-occlusion (Figure 7B). As in monocytes isolated from LPS-treated mice, monocytes exposed *in vitro* to LPS for 2 hours showed increased *IL10* and *Nos2* expression (Figure 7D). To test whether the protective activity could also be elicited by human monocytes, we exposed purified human peripheral blood monocytes to saline or LPS for 2 hours and transferred them into mice that underwent stroke as described above. Transfer of human monocytes that were exposed to LPS, but not saline, significantly reduced infarct volumes after MCAo (Figure 7C). The degree of reduction was comparable to that observed in mice receiving murine LPS-treated monocytes (-45±10% vs. -55±10%; compared to mice receiving either human or mouse PBS monocytes; P>0.05).

**Figure 7.**
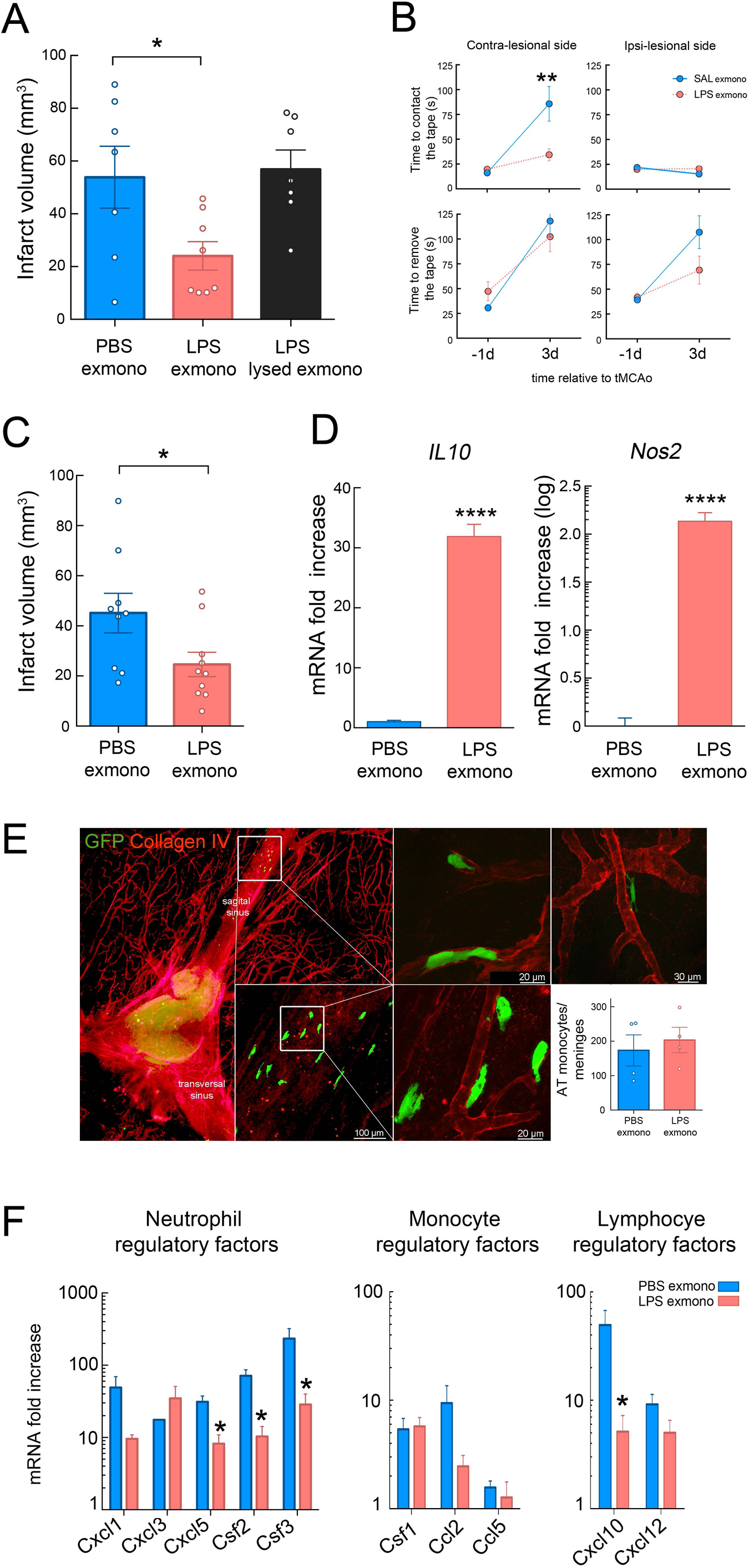
Adoptive transfer of *ex vivo* LPS-primed mouse and human monocytes exerts protection in recipient mice after MCAo. **A.** Infarct volume analysis in mice that adoptively received *ex vivo* PBS-treated monocytes (PBS exmono), LPS-treated (LPS; 100 ng/ml, 2 hours) monocytes (LPS exmonno) or lysed LPS-treated monocytes 7 hours after MCAo (LPS lysed exmono). Mice receiving *ex vivo* LPS-primed monocytes showed reduction of infarct volume (*n*=7-8 mice/group). Statistical analysis: one-way ANOVA F (2, 19) = 4.933, P=0.0188) followed by Bonferroni’s post-hoc test; *P < 0.05. **B**. Assessment of sensorimotor function by tape test performance revealed that mice receiving *ex vivo* LPS-primed monocytes had reduced latency to contact the tape on the impaired paw (contra-lesional side) than mice receiving PBS-treated monocytes at 3 days post-MCAo (*n*=13 mice/group). Statistical analysis: two-way ANOVA (interaction, F (1, 49) = 7.828, P = 0.0073; treatment, F (1, 49) = 5.915, P = 0.0187; time, F (1, 49) = 18.4, P < 0.0001) followed by Sidak’s post-hoc test; **P < 0.01. **C**. Infarct volume analysis in stroke mice receiving *ex vivo* PBS or LPS-treated human monocytes 7 hours after MCAo. LPS-primed human monocytes significantly reduced infarct volumes after MCAo when compared to mice receiving PBS-primed human monocytes (*n*=9-10 mice/group). Statistical analysis: unpaired two-tailed Student’s t test (t=2.267, df=17, *P = 0.0367). **D.** Quantification of *IL-10* and *Nos2* (iNOS) expression by qRT-PCR of *ex vivo* PBS or LPS-treated monocytes showed upregulation of both genes 2 hours after LPS treatment (*n*=4/group). Statistical analysis: unpaired two-tailed Student’s t test, *IL-10* (t=15.19, df=6, **** P < 0.0001) and *Nos2.* (t=16, df=5, **** P < 0.0001). **E**. Histology of the meninges after adoptive transfer of GFP^+^ monocytes *ex vivo* treated with LPS. Monocytes transferred 7 hours after MCAo were recruited to the meninges where they associated with vessels (extravascular and intravascular localization) identified by collagen IV expression. The graph shows the number of adoptively transferred monocytes (AT) that accumulated in the meninges at 48 hours MCAo of mice that received either PBS- or LPS-*ex vivo* treated GFP^+^ monocytes. No difference in the number of accumulated monocytes was found between groups (*n*=4 mice/group). Statistical analysis: unpaired two-tailed Student’s t test (t=0.5148, df=6, P = 0.6251). **F**. Gene expression of inflammatory molecules in the meninges of mice receiving *ex vivo* PBS-or LPS-primed monocytes 48 hours after MCAo. Decreased *Cxcl5, Csf2, Csf3* and *Cxcl10* gene upregulation was observed in the meninges of mice receiving LPS-primed monocytes (*n*=10-11 mice/group). Statistical analysis: unpaired two-tailed Student’s t test, *Cxcl5*, (t=3.6298, df=7, *P = 0.0084); *Csf2*, (t=4.4886, df=7, *P = 0.0028); *Csf3*, (t=2.6324, df=7, *P = 0.0337) and *Cxcl10*, (t=2.7547, df=7, *P = 0.0283)

We also sought to determine whether adoptively transferred monocytes exposed to LPS *ex vivo* were recruited to the brain and meninges. BM monocytes from GFP transgenic mice were exposed to vehicle (PBS) or LPS and transferred into mice 7 hours after MCAo. We found that both PBS and LPS-primed monocytes were present in the meninges but not in the brain after stroke (Figure 7E). Transferred-monocytes were mainly seen intravascularly or perivascularly in vessels in or near the superior sagittal sinus. These findings suggest that the presence of monocytes in the meninges is not sufficient to confer neuroprotection, but that priming of the monocytes by LPS is also needed.

### LPS primed monocytes suppress inflammatory cytokines in the meninges

To address whether adoptive transfer of protective monocytes would alter the inflammatory response in meninges after stroke, we conducted mRNA analysis of inflammatory molecules involved in leukocyte chemotaxis (*Ccl2, Ccl5, Cxcl1, Cxcl3, Cxcl5, Cxcl10, Cxcl12)* and survival/activation (*Csf1, Csf2, Csf3)* in the meninges of mice receiving LPS-primed monocytes 48 hours after MCAo. All inflammatory molecules were upregulated in the meninges, except for *Ccl5* (Figure 7F). Notably, neutrophil regulatory factors *Cxcl5, Csf2 and Csf3* and the lymphocyte regulatory factor *Cxcl10* were significantly decreased in mice receiving LPS-primed monocytes. These data indicate that LPS-primed monocytes in the meninges suppress the post-ischemic expression of inflammatory cytokines involved in leukocyte trafficking to the brain.

## Discussion

We have found that LPS PC leads to selective accumulation of “inflammatory” monocytes in the brains and meninges of preconditioned animals and that these monocytes are sufficient to induce neuroprotection after adoptive transfer. While previous studies have focused on the effects of LPS PC on gene regulation within the brain (Stevens et al., 2011), here we asked whether LPS PC induces a peripheral immune response that contributes to cerebral ischemic tolerance and modifies stroke outcome. We found that monocytes are recruited to brain and meninges after LPS PC. This was specific for monocytes, because neutrophils did not accumulate in the brain or meninges after LPS PC even though their frequency was strongly increased in the peripheral blood.

Importantly, we observed that monocytes isolated from the BM of mice that underwent LPS PC were able to confer robust neuroprotection when adoptively transferred to naïve mice 7 hours after MCAo, indicating that monocytes are sufficient to induce ischemic tolerance. Adoptive transfer of monocytes isolated from saline treated animals did not alter stroke outcome suggesting that LPS-induced gene expression is required to establish the protective phenotype. In contrast, adoptively transferred neutrophils isolated from the LPS-preconditioned animals or *in vitro* differentiated CAM, AAM, and MDSC were not able to confer neuroprotection. Therefore, the capacity to confer protection from ischemic brain injury is restricted to a selected monocyte subpopulation.

Our data also show that the protective phenotype of adoptively transferred monocytes is dependent on the expression of CCR2, IL-10 and iNOS. CCR2 is the major chemokine receptor involved in monocyte infiltration after brain ischemia (Gliem et al., 2012; Garcia-Bonilla et al., 2016). Here, we show that CCR2 is also required for meningeal recruitment of monocytes. Interestingly, several studies have reported increased injury after MCAo in CCR2^−^/– mice, which lack circulating inflammatory monocytes (Gliem et al., 2012; Chu et al., 2015; Garcia-Bonilla et al., 2016; Wattananit et al., 2016), an observation consistent with a neuroprotective activity of monocytes recruited to the ischemic lesion. Furthermore, hypoxic and ischemic preconditioning are also dependent on CCL2/CCR2 signaling (Rehni and Singh, 2012; Stowe et al., 2012), indicating that recruitment of protective monocytes might be a global mechanism underlying brain PC. IL-10 has been proposed as a necessary component of ischemic tolerance in the myocardium (Cai et al., 2012) and is up-regulated in the plasma, but not the brain, after LPS preconditioning (Vartanian et al., 2011). These results are consistent with a role for IL-10 production in circulating immune cells, possibly monocytes, as observed in our study. Previous studies have identified iNOS-derived NO as an essential mediator of ischemic and LPS PC (Kawano et al., 2007; Kunz et al., 2007). Our results support these previous findings and support the hypothesis that monocyte derived NO contributes to ischemic tolerance in LPS PC.

Using labeling of splenic monocyte/macrophages by subcapsular latex beads injection, we also found that LPS treatment mobilizes monocytes from the spleen into circulation and that splenic monocytes became associated with brain and meninges. The contribution of splenic monocytes to stroke pathology has been a matter of debate. While several studies have shown a role for the spleen, and specifically splenic monocytes, in promoting ischemic brain injury (Ajmo et al., 2008; Ostrowski et al., 2012; Seifert et al., 2012; Dotson et al., 2014), others failed to do so (Kim *et al.*, 2014; Zierath *et al.*, 2017). We did not specifically address the deleterious effects of splenic monocytes in brain ischemia, but our study suggests that splenic monocytes can be protective when exposed to a systemic PC stimulus. Whether this protective phenotype is also elicited during the systemic inflammatory response seen after ischemic brain injury remains to be determined.

Our study also identifies the meninges as a key immunoregulatory compartment involved in PC and ischemic brain injury. Whereas protective monocytes were found in both brain and meninges after LPS PC, they were only found in the meninges after adoptive transfer. These findings emphasize a crucial role for the meninges in mediating the neuroprotection afforded by LPS-primed monocytes. We have previously shown that meningeal γΔT cells orchestrate immune cell trafficking after ischemic brain injury by promoting neutrophil infiltration into the affected brain parenchyma (Benakis et al., 2016). Here we demonstrate that meningeal monocytes are capable of dampening the post-ischemic inflammatory response by suppressing the production of neutrophil activating and chemotactic factors such as *Csf2, Csf3* and *Cxcl5*. We have previously shown that *Csf3* is an endothelial-derived cytokine responsible for neutrophil activation after stroke (Garcia-Bonilla et al., 2015). Down-regulating *Csf3* and related cytokines in the meninges may suppress the inflammatory response in neutrophils thereby limiting their ability to invade the brain parenchyma.

Another major finding of this study is that monocytes can be primed *in vitro* to express a neuroprotective phenotype. Thus, exposure of monocytes to LPS for 2 hours was sufficient to induce their protective potential. Together with the favorable therapeutic window, the quick induction of the neuroprotective phenotype should allow for the development of clinical protocols for stroke therapy. BM mononuclear cells (BMMC) are increasingly used as a cell therapy for a variety of brain injuries including stroke, both in preclinical and clinical settings (Rosado-de-Castro et al., 2016). While the identity of the immune cells within the heterogeneous BMMC pool responsible for the protective effects has not been established, one study showed that BMMC depleted of myeloid cells lost their protective potential after transient MCAo in rats while depletion of the lymphocyte and erythrocyte lineage had no effect (Yang et al., 2016). Similarly, in rats treated with human umbilical cord blood after permanent MCAo, monocytes have been singled out as the protective cell population and peripheral blood CD14^+^ monocytes were sufficient to improve long term outcome (Womble et al., 2014). The finding that human monocytes can be reprogrammed *in vitro* to be neuroprotective is of translational and clinical relevance. While we did not address the mechanism by which human PC monocytes exert their protective effects, it is likely, that their effector mechanisms are also dependent upon IL-10 and NO. Importantly, iNOS and IL-10 are upregulated in LPS-treated human monocytes and human IL-10 shows cross-species activity on mouse cells (Tan et al., 1993).

In summary, our study identifies a distinct population of splenic monocytes as key effectors cell in the profound protective effect afforded by LPS PC. After exposure to LPS either *in vitro* or *in vivo*, these cells are able to modulate meningeal and parenchymal immune responses after MCAo, limiting ischemic injury and improving functional outcome with a clinically relevant therapeutic window. Furthermore, our findings demonstrate that *in vitro* priming of monocytes with TLR4 agonists triggers protective potential against ischemic brain injury and raise the possibility that similar approaches could be used to enhance the effectiveness of monocyte-based cell therapies in ischemic stroke patients.

## Acknowledgements

This work was supported by the National Institutes of Health NS34179 (C.I) and NS081179 (J.A) grants and by the American Heart Association Grant 16SDG30970036 (L.G). The generous support of the Feil Family Foundation is gratefully acknowledged.

## References

Ahmed SH, He YY, Nassief A, Xu J, Xu XM, Hsu CY, Faraci FM (2000) Effects of lipopolysaccharide priming on acute ischemic brain injury. Stroke 31:193–199.

Ajmo CT, Vernon DOL, Collier L, Hall AA, Garbuzova-Davis S, Willing A, Pennypacker KR (2008) The spleen contributes to stroke-induced neurodegeneration. J Neurosci Res 86:2227–2234.

Arnold L, Henry A, Poron F, Baba-Amer Y, van Rooijen N, Plonquet A, Gherardi RK, Chazaud B (2007) Inflammatory monocytes recruited after skeletal muscle injury switch into antiinflammatory macrophages to support myogenesis. J Exp Med 204:1057–1069.

Auffray C, Fogg D, Garfa M, Elain G, Join-Lambert O, Kayal S, Sarnacki S, Cumano A, Lauvau G, Geissmann F (2007) Monitoring of Blood Vessels and Tissues by a Population of Monocytes with Patrolling Behavior. Science 317:666–670.

Banks WA, Robinson SM (2010) Minimal penetration of lipopolysaccharide across the murine blood-brain barrier. Brain Behavior and Immunity 24:102–109.

Bauer P, Welbourne T, Shigematsu T, Russell J, Granger DN (2000) Endothelial expression of selectins during endotoxin preconditioning. Am J Physiol Regul Integr Comp Physiol 279:R2015–R2021.

Benakis C, Brea D, Caballero S, Faraco G, Moore J, Murphy M, Sita G, Racchumi G, Ling L, Pamer EG, Iadecola C, Anrather J (2016) Commensal microbiota affects ischemic stroke outcome by regulating intestinal γδ T cells. Nat Med 22:516–523.

Biswas SK, Lopez-Collazo E (2009) Endotoxin tolerance: new mechanisms, molecules and clinical significance. Trends in Immunology 30:475–487.

Bronte V (2009) Myeloid-derived suppressor cells in inflammation: uncovering cell subsets with enhanced immunosuppressive functions. Eur J Immunol 39:2670–2672.

Cai ZP, Parajuli N, Zheng X, Becker L (2012) Remote ischemic preconditioning confers late protection against myocardial ischemia-reperfusion injury in mice by upregulating interleukin-10. Basic Res Cardiol 107:277.

Chu HX, Broughton BRS, Ah Kim H, Lee S, Drummond GR, Sobey CG (2015) Evidence That Ly6Chi Monocytes Are Protective in Acute Ischemic Stroke by Promoting M2 Macrophage Polarization. Stroke:STROKEAHA.115.009426.

Corraliza IM, Campo ML, Soler G, Modolell M (1994) Determination of arginase activity in macrophages: a micromethod. J Immunol Methods 174:231–235.

Dotson AL, Wang J, Saugstad J, Murphy SJ, Offner H (2014) Splenectomy reduces infarct volume and neuroinflammation in male but not female mice in experimental stroke. J Neuroimmunol.

Dunning MJ, Smith ML, Ritchie ME, Tavaré S (2007) beadarray: R classes and methods for Illumina bead-based data. Bioinformatics 23:2183–2184.

Forghani P, Khorramizadeh MR, Waller EK (2012) Natural suppressor cells; past, present and future. Front Biosci (Elite Ed) 4:1237–1245.

Garcia-Bonilla L, Benakis C, Moore J, Iadecola C, Anrather J (2014a) Immune mechanisms in cerebral ischemic tolerance. Front Neurosci 8:44.

Garcia-Bonilla L, Faraco G, Moore J, Murphy M, Racchumi G, Srinivasan J, Brea D, Iadecola C, Anrather J (2016) Spatio-temporal profile, phenotypic diversity, and fate of recruited monocytes into the post-ischemic brain. Journal of Neuroinflammation 13:285.

Garcia-Bonilla L, Moore JM, Racchumi G, Zhou P, Butler JM, Iadecola C, Anrather J (2014b) Inducible nitric oxide synthase in neutrophils and endothelium contributes to ischemic brain injury in mice. The Journal of Immunology 193:2531–2537.

Garcia-Bonilla L, Racchumi G, Murphy M, Anrather J, Iadecola C (2015) Endothelial CD36 Contributes to Postischemic Brain Injury by Promoting Neutrophil Activation via CSF3. J Neurosci 35:14783–14793.

Geissmann F, Jung S, Littman DR (2003) Blood monocytes consist of two principal subsets with distinct migratory properties. Immunity.

Gliem M, Mausberg AK, Lee J-I, Simiantonakis I, van Rooijen N, Hartung H-P, Jander S (2012) Macrophages prevent hemorrhagic infarct transformation in murine stroke models. Ann Neurol 71:743–752.

Highfill SL, Rodriguez PC, Zhou Q, Goetz CA, Koehn BH, Veenstra R, Taylor PA, Panoskaltsis-Mortari A, Serody JS, Munn DH, Tolar J, Ochoa AC, Blazar BR (2010) Bone marrow myeloid-derived suppressor cells (MDSCs) inhibit graft-versus-host disease (GVHD) via an arginase-1-dependent mechanism that is up-regulated by interleukin-13. Blood 116:5738–5747.

Huang B, Pan P-Y, Li Q, Sato AI, Levy DE, Bromberg J, Divino CM, Chen S-H (2006) Gr-1+CD115+ immature myeloid suppressor cells mediate the development of tumor-induced T regulatory cells and T-cell anergy in tumor-bearing host. Cancer Res 66:1123–1131.

Iadecola C, Anrather J (2011) Stroke research at a crossroad: asking the brain for directions. 14:1363–1368.

Jackman K, Kunz A, Iadecola C (2011) Modeling focal cerebral ischemia in vivo. Methods Mol Biol 793:195–209.

Kawano T, Kunz A, Abe T, Girouard H, Anrather J, Zhou P, Iadecola C (2007) iNOS-derived NO and nox2-derived superoxide confer tolerance to excitotoxic brain injury through peroxynitrite. J Cereb Blood Flow Metab 27:1453–1462.

Kilkenny C, Browne W, Cuthill IC, Emerson M, Altman DG, National Centre for the Replacement, Refinement and Reduction of Amimals in Research (2011) Animal research: reporting in vivo experiments--the ARRIVE guidelines. Journal of Cerebral Blood Flow & Metabolism 31:991–993.

Kim E, Yang J D Beltran C, Cho S (2014) Role of spleen-derived monocytes/macrophages in acute ischemic brain injury. Journal of Cerebral Blood Flow & Metabolism.

Kirino T (2002) Ischemic tolerance. J Cereb Blood Flow Metab 22:1283–1296.

Kunz A, Park L, Abe T, Gallo EF, Anrather J, Zhou P, Iadecola C (2007) Neurovascular protection by ischemic tolerance: role of nitric oxide and reactive oxygen species. J Neurosci 27:7083–7093.

Liesz A, Mracsko E, Zhou W, Yang H, Na S-Y, Tracey KJ, Veltkamp R (2015) DAMP Signaling is a Key Pathway Inducing Immune Modulation after Brain Injury. J Neurosci 35:583–598.

Lin H-Y, Huang C-C, Chang K-F (2009) Lipopolysaccharide preconditioning reduces neuroinflammation against hypoxic ischemia and provides long-term outcome of neuroprotection in neonatal rat. Pediatr Res 66:254–259.

Livak KJ, Schmittgen TD (2001) Analysis of relative gene expression data using real-time quantitative PCR and the 2(-Delta Delta C(T)) Method. Methods 25:402–408.

Louveau A, Kipnis J (2015) Dissection and immunostaining of mouse whole-mount meninges. Protocol Exchange.

MacMicking JD, Nathan C, Hom G, Chartrain N, Fletcher DS, Trumbauer M, Stevens K, Xie QW, Sokol K, Hutchinson N (1995) Altered responses to bacterial infection and endotoxic shock in mice lacking inducible nitric oxide synthase. Cell 81:641–650.

Manthey CL, Vogel SN (1994) Elimination of trace endotoxin protein from rough chemotype LPS. Innate Immun 1:84–91.

Marsh B, Stevens SL, Packard AEB, Gopalan B, Hunter B, Leung PY, Harrington CA, Stenzel-Poore MP (2009a) Systemic lipopolysaccharide protects the brain from ischemic injury by reprogramming the response of the brain to stroke: a critical role for IRF3. J Neurosci 29:9839–9849.

Marsh BJ, Stenzel-Poore MP (2008) Toll-like receptors: novel pharmacological targets for the treatment of neurological diseases. Current Opinion in Pharmacology 8:8–13.

Marsh BJ, Williams-Karnesky RL, Stenzel-Poore MP (2009b) Toll-like receptor signaling in endogenous neuroprotection and stroke. Neuroscience 158:1007–1020.

Miró-Mur F, Pérez-de Puig I, Ferrer-Ferrer M, Urra X, Justicia C, Chamorro A, Planas AM (2015) Immature monocytes recruited to the ischemic mouse brain differentiate into macrophages with features of alternative activation. Brain Behavior and Immunity 53:18–33.

Morham SG, Langenbach R, Loftin CD, Tiano HF, Vouloumanos N, Jennette JC, Mahler JF, Kluckman KD, Ledford A, Lee CA, Smithies O (1995) Prostaglandin synthase 2 gene disruption causes severe renal pathology in the mouse. Cell 83:473–482.

Muzio M, Bosisio D, Polentarutti N, D’amico G, Stoppacciaro A, Mancinelli R, van’t Veer C, Penton-Rol G, Ruco LP, Allavena P, Mantovani A (2000) Differential expression and regulation of toll-like receptors (TLR) in human leukocytes: selective expression of TLR3 in dendritic cells. J Immunol 164:5998–6004.

Nahrendorf M, Pittet MJ, Swirski FK (2010) Monocytes: Protagonists of Infarct Inflammation and Repair After Myocardial Infarction. Circulation 121:2437–2445.

Nahrendorf M, Swirski FK, Aikawa E, Stangenberg L, Wurdinger T, Figueiredo J-L, Libby P, Weissleder R, Pittet MJ (2007) The healing myocardium sequentially mobilizes two monocyte subsets with divergent and complementary functions. J Exp Med 204:3037–3047.

Ostrand-Rosenberg S (2010) Myeloid-derived suppressor cells: more mechanisms for inhibiting antitumor immunity. Cancer Immunol Immunother 59:1593–1600.

Ostrowski RP, Schulte RW, Nie Y, Ling T, Lee T, Manaenko A, Gridley DS, Zhang JH (2012) Acute splenic irradiation reduces brain injury in the rat focal ischemic stroke model. Transl Stroke Res 3:473–481.

Pradillo JM, Fernández-López D, García-Yébenes I, Sobrado M, Hurtado O, Moro MA, Lizasoain I (2009) Toll-like receptor 4 is involved in neuroprotection afforded by ischemic preconditioning. J Neurochem 109:287–294.

Rehni AK, Singh TG (2012) Involvement of CCR-2 chemokine receptor activation in ischemic preconditioning and postconditioning of brain in mice. Cytokine 60:83–89.

Reich M, Liefeld T, Gould J, Lerner J, Tamayo P, Mesirov JP (2006) GenePattern 2.0. Nat Genet 38:500–501.

Ritchie ME, Phipson B, Wu D, Hu Y, Law CW, Shi W, Smyth GK (2015) limma powers differential expression analyses for RNA-sequencing and microarray studies. Nucleic Acids Res 43:e47–e47.

Rosado-de-Castro PH, de Carvalho FG, de Freitas GR, Mendez-Otero R, Pimentel-Coelho PM (2016) Review of Preclinical and Clinical Studies of Bone Marrow-Derived Cell Therapies for Intracerebral Hemorrhage. Stem Cells Int 2016:4617983–18.

Rosenzweig HL, Lessov NS, Henshall DC, Minami M, Simon RP, Stenzel-Poore MP (2004) Endotoxin preconditioning prevents cellular inflammatory response during ischemic neuroprotection in mice. Stroke 35:2576–2581.

Rosenzweig HL, Minami M, Lessov NS, Coste SC, Stevens SL, Henshall DC, Meller R, Simon RP, Stenzel-Poore MP (2007) Endotoxin preconditioning protects against the cytotoxic effects of TNFalpha after stroke: a novel role for TNFalpha in LPS-ischemic tolerance. J Cereb Blood Flow Metab 27:1663–1674.

Seifert HA, Leonardo CC, Hall AA, Rowe DD, Collier LA, Benkovic SA, Willing AE, Pennypacker KR (2012) The spleen contributes to stroke induced neurodegeneration through interferon gamma signaling. Metab Brain Dis 27:131–141.

Serafini P, De Santo C, Marigo I, Cingarlini S, Dolcetti L, Gallina G, Zanovello P, Bronte V (2004) Derangement of immune responses by myeloid suppressor cells. Cancer Immunol Immunother 53:64–72.

Serbina NV, Jia T, Hohl TM, Pamer EG (2008) Monocyte-mediated defense against microbial pathogens. Annu Rev Immunol 26:421–452.

Sherman BT, Huang DW, Tan Q, Guo Y, Bour S, Liu D, Stephens R, Baseler MW, Lane HC, Lempicki RA (2007) DAVID Knowledgebase: a gene-centered database integrating heterogeneous gene annotation resources to facilitate high-throughput gene functional analysis. BMC Bioinformatics 8:426.

Shi C, Jia T, Mendez-Ferrer S, Hohl TM, Serbina NV, Lipuma L, Leiner I, Li MO, Frenette PS, Pamer EG (2011) Bone marrow mesenchymal stem and progenitor cells induce monocyte emigration in response to circulating toll-like receptor ligands. Immunity 34:590–601.

Singh AK, Jiang Y (2004) How does peripheral lipopolysaccharide induce gene expression in the brain of rats? Toxicology 201:197–207.

Smith RM, Lecour S, Sack MN (2002) Innate immunity and cardiac preconditioning: a putative intrinsic cardioprotective program. Cardiovasc Res 55:474–482.

Stevens SL, Leung PY, Vartanian KB, Gopalan B, Yang T, Simon RP, Stenzel-Poore MP (2011) Multiple preconditioning paradigms converge on interferon regulatory factor-dependent signaling to promote tolerance to ischemic brain injury. J Neurosci 31:8456–8463.

Stowe AM, Wacker BK, Cravens PD, Perfater JL, Li MK, Hu R, Freie AB, Stüve O, Gidday JM (2012) CCL2 upregulation triggers hypoxic preconditioning-induced protection from stroke. Journal of Neuroinflammation 9:33.

Swirski FK, Libby P, Aikawa E, Alcaide P, Luscinskas FW, Weissleder R, Pittet MJ (2007) Ly-6Chi monocytes dominate hypercholesterolemia-associated monocytosis and give rise to macrophages in atheromata. J Clin Invest 117:195–205.

Swirski FK, Nahrendorf M, Etzrodt M, Wildgruber M, Cortez-Retamozo V, Panizzi P, Figueiredo J-L, Kohler RH, Chudnovskiy A, Waterman P, Aikawa E, Mempel TR, Libby P, Weissleder R, Pittet MJ (2009) Identification of splenic reservoir monocytes and their deployment to inflammatory sites. Science 325:612–616.

Swirski FK, Wildgruber M, Ueno T, Figueiredo J-L, Panizzi P, Iwamoto Y, Zhang E, Stone JR, Rodriguez E, Chen JW, Pittet MJ, Weissleder R, Nahrendorf M (2010) Myeloperoxidase-rich Ly-6C+ myeloid cells infiltrate allografts and contribute to an imaging signature of organ rejection in mice. J Clin Invest 120:2627–2634.

Tacke F, Alvarez D, Kaplan TJ, Jakubzick C, Spanbroek R, Llodra J, Garin A, Liu J, Mack M, van Rooijen N, Lira SA, Habenicht AJ, Randolph GJ (2007) Monocyte subsets differentially employ CCR2, CCR5, and CX3CR1 to accumulate within atherosclerotic plaques. J Clin Invest 117:185–194.

Tan JC, Indelicato SR, Narula SK, Zavodny PJ, Chou CC (1993) Characterization of interleukin-10 receptors on human and mouse cells. Journal of Biological Chemistry 268:21053–21059.

Tasaki K, Ruetzler CA, Ohtsuki T, Martin D, Nawashiro H, Hallenbeck JM (1997) Lipopolysaccharide pre-treatment induces resistance against subsequent focal cerebral ischemic damage in spontaneously hypertensive rats. Brain Res 748:267–270.

Vartanian KB, Stevens SL, Marsh BJ, Williams-Karnesky R, Lessov NS, Stenzel-Poore MP (2011) LPS preconditioning redirects TLR signaling following stroke: TRIF-IRF3 plays a seminal role in mediating tolerance to ischemic injury. Journal of Neuroinflammation 8:140.

Wang F, Birch SE, He R, Tawadros P, Szaszi K, Kapus A, Rotstein OD (2010) Remote Ischemic Preconditioning by Hindlimb Occlusion Prevents Liver Ischemic/Reperfusion Injury. Ann Surg 251:292–299.

Wattananit S, Tornero D, Graubardt N, Memanishvili T, Monni E, Tatarishvili J, Miskinyte G, Ge R, Ahlenius H, Lindvall O, Schwartz M, Kokaia Z (2016) Monocyte-Derived Macrophages Contribute to Spontaneous Long-Term Functional Recovery after Stroke in Mice. J Neurosci 36:4182–4195.

Weischenfeldt J, Porse B (2008) Bone Marrow-Derived Macrophages (BMM): Isolation and Applications. CSH Protoc 2008:pdb.prot5080–pdb.prot5080.

Womble TA, Green S, Shahaduzzaman M, Grieco J, Sanberg PR, Pennypacker KR, Willing AE (2014) Monocytes are essential for the neuroprotective effect of human cord blood cells following middle cerebral artery occlusion in rat. Mol Cell Neurosci 59:76–84.

Yang B, Parsha K, Schaar K, Xi X, Aronowski J, Savitz SI (2016) Various Cell Populations Within the Mononuclear Fraction of Bone Marrow Contribute to the Beneficial Effects of Autologous Bone Marrow Cell Therapy in a Rodent Stroke Model. Transl Stroke Res:1–9.

Youn J-I, Nagaraj S, Collazo M, Gabrilovich DI (2008) Subsets of myeloid-derived suppressor cells in tumor-bearing mice. The Journal of Immunology 181:5791–5802.

Zierath D, Shen A, Stults A, Olmstead T, Becker KJ (2017) Splenectomy Does Not Improve Long-Term Outcome After Stroke. Stroke:STROKEAHA.116.016037.

